# Sample-derived cDNA guides broad host RNA depletion for *in vivo* pathogen transcriptomics

**DOI:** 10.64898/2026.03.12.711445

**Authors:** Tugrul Doruk, Oytun Sarigöz, Ummehan Avican, Rikki Franklin Fredriksen, Leyden Fernandez, Kemal Avican

## Abstract

Profiling bacterial transcriptomes during active infection is fundamentally limited by the overwhelming dominance of host RNA, which reduces pathogen-derived reads to a fraction of the total sequencing output. Standard ribosomal RNA depletion kits remove rRNA but leave a vast pool of host messenger and non-coding RNAs intact, necessitating prohibitively deep sequencing to achieve adequate bacterial coverage. Here, we present a sample-derived, cDNA-guided broad host RNA depletion method that selectively eliminates host transcripts directly from total host–microbe RNA mixtures. By reverse-transcribing host RNA into cDNA and using it to guide RNase H-mediated cleavage of host RNA:cDNA duplexes, our approach depletes a broad spectrum of host RNA biotypes including rRNA, non-coding RNA, and mRNA without requiring custom probe design. Validated in both mouse and human RNA backgrounds using defined 1% bacterial spike-ins, our method enriches bacterial reads over 14-fold and outperforms commercial alternatives. Supplemental oligo-probes targeting highly structured 7S RNA and 18S rRNA species, which are refractory to cDNA synthesis, further improve depletion efficiency. Critically, depletion does not distort bacterial transcript coverage, expression distributions, or differential expression profiles. Rarefaction analysis demonstrates that fully saturated bacterial transcriptome coverage is achievable at 50–100 million reads in enriched samples, compared to over 600 million reads in non-depleted libraries. Applied to *in vivo Salmonella enterica* serovar Typhimurium infection models spanning acute and chronic disease stages, cDNA-guided depletion consistently enriched bacterial reads while preserving transcriptome-wide concordance with non-depleted samples and improving detection of differentially expressed genes. This methodology offers a robust, cost-effective, and broadly applicable solution for bacterial transcriptomics directly from infected tissues.

## Introduction

Deciphering microbial gene expression is fundamental to understanding host–microbe interactions, community dynamics, and pathogen biology. Over the past two decades, rapid advances in next generation sequencing technologies have transformed microbial transcriptomics, enabling large-scale detection and quantification of gene expression within microbial populations, and across complex communities^1^. These technologies are distinguished by their ability to accurately quantify gene expression with exceptional sensitivity and specificity across both eukaryotic and prokaryotic systems, yielding unprecedented insights into microbial biology, community dynamics, and host–microbe interactions^2–6^. Among them, RNA sequencing (RNA-seq) has emerged as a cornerstone approach, generating millions to billions of reads per run to enable high-resolution transcript profiling. Despite its power, RNA-seq faces a critical barrier: the overwhelming dominance of ribosomal RNA (rRNA), which typically accounts for >80% of total RNA, leaving only a small fraction as more informative messenger RNA (mRNA) and regulatory RNAs^7–10^. Two strategies are commonly employed to mitigate this challenge: (i) increasing sequencing depth and (ii) enriching informative transcripts by depleting abundant RNA species^11^. Depletion of unwanted RNA species increases the proportion of RNA sequenced, which can significantly decrease sequencing costs, especially in cases where host RNA vastly outnumbers bacterial RNA.

Ribosomal RNA depletion from a total RNA sample has become standard practice for RNA-seq experiments focused predominantly on mRNA^10,12,13^. Several methods are effectively employed for rRNA depletion, though their efficiency varies with sample type and composition^14^. A commonly used approach relies on hybridization of rRNA-specific oligonucleotides to target rRNA molecules, followed by capture and removal with magnetic beads^15,16^. Another method is based on enzymatic digestion of the rRNA species after hybridization with antisense oligonucleotide probes using nucleases such as RNase H^17^. Generally, these depletion strategies typically employ oligonucleotides that are complementary to rRNA sequences, allowing their selective removal from total RNA samples. Commercial kits implementing these approaches are widely available, and numerous studies have compared their efficiencies^10,14,18–20^. Yet, these kits frequently underperform when rRNA sequences diverge from probe designs or when RNA integrity is compromised^20^. Moreover, their scalability is constrained by input requirements and concentration thresholds (typically 10–100 ng of total RNA), limiting cost-effectiveness for bacterial transcriptomics^21^. These limitations of commercial kits led to development of alternative methods for rRNA depletion. For example, Armour, et al.^22^ developed a method in which the use of a non-random primer mixture during the cDNA synthesis step effectively depletes rRNA species prior to RNA-seq library preparation^22^. Huang, et al.^23^ performed rRNA depletion using probes generated through a computational tool^23^. Betin, et al.^21^ developed a hybridization-based capture method (PatH-Cap) to selectively enrich pathogen mRNA from mixed host–pathogen samples, thereby enhancing bacterial transcript detection and enabling more effective dual RNA-seq analysis^21^. Wangsanuwat, et al.^24^ developed EMBR-seq, a cost-effective rRNA depletion strategy that employs blocking primers at rRNA termini to prevent their cDNA synthesis, enabling efficient bacterial mRNA enrichment even from ultra-low RNA inputs^24^. Choe, et al.^25^ introduced RiboRid, a cost-effective and highly efficient rRNA depletion method for bacterial transcriptomics^25^. Recently, Qasim and Sarin^26^ developed DepStep, a one-step rRNA depletion method that applies species-specific biotinylated antisense probes to selectively bind and eliminate target rRNA molecules^26^.

While the current strategies effectively address rRNA dominance in pure or enriched microbial cultures, transcriptomic profiling of bacteria directly within host tissues during active infection presents a more complex analytical challenge. In *in vivo* infection models and clinical biopsy specimens, invading bacteria are vastly outnumbered by host cells, resulting in remarkably low abundance of bacterial RNA within total RNA extracts^27^. The consequent co-extraction of host RNA at overwhelming excess constitutes a severe technical bottleneck for downstream sequencing applications, as host transcripts consume sequencing capacity and reduce coverage of the bacterial genome to levels insufficient for meaningful interpretation. It is estimated that capturing both host and pathogen transcripts simultaneously requires approximately 2 billion reads from total RNA libraries, or 200 million reads from rRNA-depleted samples^28^. Under these *in vivo* conditions, standard commercial rRNA depletion kits prove largely inadequate^29^. Although such kits successfully remove ribosomal sequences, they leave behind a vast pool of host messenger and non-coding RNAs that continue to outcompete the rare bacterial transcripts for sequencing reads. To achieve the depth of coverage required for cost-effective and comprehensive analysis of pathogen gene expression *in vivo,* innovative depletion strategies are required to bridge the gap between sequencing cost and data resolution.

The central objective of available approaches is to maximize the bacteria-to-host RNA ratio while preserving the true physiological proportions of individual bacterial transcripts as they exist within infected tissue. Several methodological solutions have been described to address this challenge. Differential lysis protocols exploit intrinsic structural differences between host and bacterial cell membranes to selectively rupture eukaryotic cells and degrade host nucleic acids prior to bacterial RNA extraction, substantially reducing host-derived contamination^30^. More advanced methodologies circumvent conventional depletion entirely through cDNA-RNA subtractive hybridization or the physical enrichment based on fluorescent sorting and antibody based selection of intact bacteria from host tissue prior to lysis^31^. Despite continued methodological progress, removal of host RNA background to obtain fully representative bacterial transcriptomes remains a defining challenge in elucidating the molecular mechanisms of bacterial pathogenesis and host-microbe interaction *in vivo*.

To overcome the overwhelming host RNA background in infected tissue extracts, we developed a broadly applicable method that selectively eliminates host-derived transcripts from total host-microbe RNA mixtures, enabling comprehensive profiling of the bacterial transcriptome. Our method generates cDNAs complementary to host RNA species; upon hybridization, the resulting cDNA:RNA duplexes are selectively cleaved by RNase H, enabling efficient and specific enrichment of target transcripts. Unlike conventional approaches, our strategy leverages cDNAs derived from the sample itself as depletion probes. This approach is highly scalable, maintaining efficiency across a wide range of RNA input concentrations without the performance constraints observed in commercial kits. Applied to infected tissue samples, our method substantially reduces sequencing depth requirements while delivering comprehensive pathogen transcriptome coverage, offering a robust and cost-effective alternative for bacterial transcriptomics.

## Results

### Enrichment of bacterial RNA with cDNA-guided depletion of host RNAs

A single bacterial cell carries ∼100 fg of total RNA, whereas a typical mammalian cell contains ∼30,000 fg, ∼300× more than bacterial cell (**Fig. 1a**). As a result, even when host and pathogen cell numbers are equal, bacterial transcripts make up only a small fraction of total RNA and can drop by orders of magnitude *in vivo*. Without depletion, achieving adequate coverage of bacterial transcripts would require billions of reads per sample; in our schematic, ∼3–9 billion reads are needed to obtain comparable information content (**Fig. 1a**). Depleting only host rRNA reduces this burden but still leaves most host mRNA/tRNA intact, yielding at best a modest enrichment and requiring ∼600–1,800 million reads. In contrast, broad host total RNA depletion (almost all types) would substantially enrich bacterial RNA and lower sequencing depth to ∼10–30 million reads (**Fig. 1a**). To achieve this, we optimized a cDNA-guided RNase H workflow that selectively eliminates host RNAs directly from total RNA (**Fig. 1b**). Briefly, cDNA is synthesized with random hexamers from either uninfected host RNA (host-only) or infected tissue RNA (mixed RNA). These cDNAs are hybridized against total RNA from infected tissue, allowing RNase H to cleave the resulting cDNA:RNA duplexes. A subsequent DNase step removes the cDNA, leaving behind an RNA pool enriched for bacterial transcripts. (**Fig. 1b**). Using cDNA derived from the infected sample itself adapts the probe set to the sample’s host transcriptome without custom design, while the rarity of pathogen transcripts minimizes the chance of off-target depletion of bacterial RNAs).

**Fig. 1.**
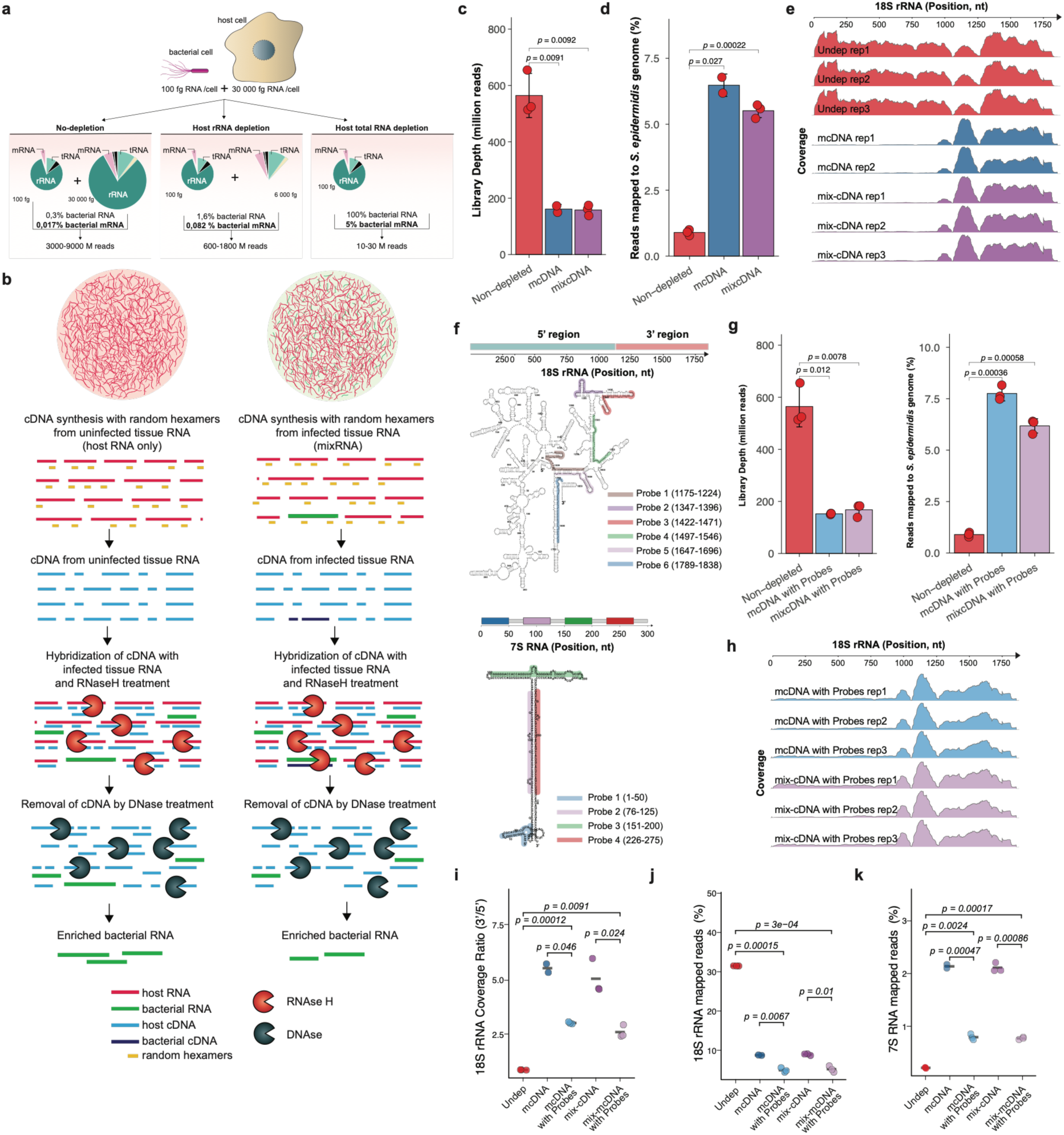
Enrichment strategy and performance of cDNA-guided host RNA depletion in mouse RNA mixtures. **(a)** Expected sequencing burden for bacterial *in vivo* RNA-seq for a 1:1 ratio of host-to-pathogen cells. Conceptual comparison of host versus bacterial RNA abundance per cell (100 fg vs. 30,000 fg) and the implied sequencing depth required for non-depleted (∼3–9 billion reads), host rRNA-only depleted (∼70–210 million reads), and host total RNA depleted (∼10–30 million reads) samples. **(b)** Workflow: cDNA generation with random hexamers from uninfected or infected tissue RNA; hybridization to total RNA from infected tissue; RNase H digestion of cDNA:RNA hybrids; DNase cleanup; enriched bacterial RNA. Red and green lines represent host RNA and bacterial RNA, respectively. Short gray lines indicate random hexamers used in cDNA synthesis, and blue lines indicate cDNA synthesized from host and bacterial RNA. **(c)** Overview of experimental design for mouse total RNA mixed with 1% *S. epidermidis* RNA. **(d)** Fraction of reads mapping to *S. epidermidis* in mouse RNA mixtures after depletion with mcDNA or mix-mcDNA. **(e)** Read depth across the mouse 18S rRNA locus (1,869 nt) in non-depleted (red), mcDNA-depleted (blue), and mix-mcDNA-depleted (purple) samples. **(f)** Secondary structure models of mouse 18S rRNA (top) and 7S RNA (bottom) with the positions of supplemental oligo-probes indicated by colored highlights. For 18S rRNA, six probes (Probes 1–6) target the 3’ region (positions 1,175–1,838 nt). For 7S RNA, four probes (Probes 1–4) are distributed across the full length of the transcript (positions 1–275 nt). Probe positions are mapped onto both the linear sequence (color-coded bars) and the secondary structure. **(g)** Bar plots showing library sequencing depth (left, million reads) and the percentage of reads mapping to the *S. epidermidis* genome (right) for non-depleted, mcDNA-depleted with probes, and mix-mcDNA-depleted with probes samples. **(h)** Read depth across the mouse 18S rRNA locus (1,869 nt) in mcDNA-depleted (blue) and mix-mcDNA-depleted (purple) samples supplemented with targeted oligo-probes across biological replicates. **(i)** 18S rRNA 3’/5’ coverage ratio across depletion conditions. Both mcDNA and mix-mcDNA depletion without probes showed significantly elevated 3’/5’ ratios compared to non-depleted samples (p=0.00012 and p=0.046, respectively). Probe supplementation significantly reduced this imbalance in both conditions (mcDNA with probes: p=0.0091; mix-mcDNA with probes: p=0.024). **(j)** Percentage of reads mapping to 18S rRNA. Probe-supplemented depletion significantly reduced 18S rRNA reads compared to both non-depleted samples (p=3×10⁻⁴ and p=0.00015) and probe-free depleted conditions (mcDNA: p=0.0067; mix-mcDNA: p=0.01). **(k)** Percentage of reads mapping to 7S RNA. Probe supplementation significantly reduced 7S RNA reads across all comparisons (p-values ranging from 0.00017 to 0.00086). Data represent individual replicates; horizontal bars indicate group medians. FDR-adjusted Welch’s t-test. n = 3 biological replicates for panels (c), (d), (g), (i), (j) and (k), except for mcDNA-depleted samples, here n = 2 owing to incomplete library preparation.

### cDNA-guided depletion of host RNAs enriches bacterial RNA

To optimize our depletion approach, we generated mixed RNA samples by spiking 1% *Staphylococcus epidermidis* total RNA into mouse total RNA, using *S. epidermidis* as a representative Gram-positive pathogen. With these samples we varied the reverse transcriptase, the RNase H enzyme, its concentration and treatment time, the concentration and source of cDNA, the bacterial RNA content of the target sample, and the probe concentration (Supplementary Note S1 and Supplementary Fig. S1). To test whether the cDNA source influences depletion efficiency, cDNAs were synthesized either from mouse host RNA alone (mcDNA), prepared from FVB/N mouse mesenteric lymph nodes, or from the 1% bacterial-mouse RNA mixture itself (mix-mcDNA). These cDNAs were hybridized to fresh 1% bacterial-mouse RNA mixtures, the resulting RNA:DNA hybrids were digested with RNase H, and residual cDNA was removed by DNase treatment. The depleted RNA, together with non-depleted 1% bacterial-mouse control mixtures, was used for RNA-seq library preparation, yielding comparable insert sizes across all libraries (**Supplementary Fig. S2**). To determine the sequencing depth required for full bacterial transcriptome coverage, libraries were initially sequenced shallowly and then extended to up to 650 million reads for non-depleted samples; bacterial-only control libraries were sequenced at lower depths because they saturate earlier **(Fig. 1c).**

Non-depleted samples yielded ∼1% of reads mapping to the *S. epidermidis* genome, consistent with the input mixing ratio (**Fig. 1d**). Host RNA depletion significantly improved bacterial read recovery: mcDNA depletion yielded up to 7% bacterial reads and mix-mcDNA yielded 5.5% **(Fig. 1d)**. Inspecting coverage of the most abundant host RNA species revealed that the 3’ region of 18S rRNA was not depleted as efficiently as its 5’ region (**Fig. 1e**), and 7S RNA was likewise under-depleted across its full length. Both RNAs are characterized by extensive secondary structure (**Fig. 1f**), which we reasoned might impair efficient cDNA synthesis by random hexamer priming, resulting in poor cDNA representation of these regions and consequently inadequate depletion. To address this, we designed antisense oligonucleotide probes covering the entire mouse 7S RNA and the 3’ region of 18S rRNA (**Fig. 1f, Supplementary Table S1**) and supplemented the cDNA bait pool with these probes in subsequent depletion reactions. With lower sequencing depth than the non-depleted samples (**Fig. 1g**), probe supplementation further increased bacterial RNA enrichment, to up to 8% with mcDNA and 6.5% with mix-mcDNA (**Fig. 1g**). The prominent 3’ residual coverage peak observed in probe-free conditions was substantially reduced in both mcDNA- and mix-mcDNA-depleted samples across all biological replicates (**Fig. 1h**). Probe supplementation also significantly increased the 3’/5’ coverage ratio of 18S rRNA reads and reduced the overall fraction of reads mapping to 18S rRNA compared with unsupplemented depletion (**Fig. 1i and j**), indicating that the probes effectively patched the previously under-depleted 3’ region and depleted more 18S rRNA molecules. Similarly, 7S RNA probe supplementation reduced the fraction of reads mapping to 7S RNA, confirming improved depletion of this species (**Fig. 1k**).

### cDNA-guided depletion outperforms commercial enrichment tools

Having validated the cDNA-guided depletion approach in mouse RNA, we next extended this strategy to human RNA samples. We also compared our method with two commonly used enrichment strategies: RiboZero Gold, which depletes both host and bacterial rRNA, and MICROB*Enrich*, which targets multiple host rRNAs and mRNAs to enrich for bacterial transcripts. As with the mouse experiments, cDNA was generated from either human host RNA alone (hcDNA), extracted from Caco-2 cells, or from a 1% bacterial–human RNA mixture (mix-hcDNA). Given the high sequence and structure conservation of 7S RNA and 18S rRNA in both mouse and human (**Supplementary Fig. S3**), the same supplemental oligo-probes used in the mouse experiments were retained to ensure adequate depletion coverage of these highly structured RNA species. The same tiered sequencing depth strategy employed for mouse samples was applied here, with non-depleted human libraries sequenced up to 650 million reads (**Fig. 2a**) Consistent with the mouse results, non-depleted samples yielded approximately 1% of reads mapping to the *S. epidermidis* genome, reflecting the expected mixing ratio (**Fig. 2b**). MICROB*Enrich* significantly increased bacterial read recovery to approximately 7-fold over non-depleted samples, whereas RiboZero Gold achieved only ∼2-fold enrichment. The modest performance of RiboZero Gold likely reflects its co-depletion of bacterial rRNA alongside host rRNA, which reduces overall bacterial transcript recovery. In contrast, our cDNA-guided depletion markedly outperformed both commercial methods, enriching bacterial RNA reads to over 14-fold with hcDNA and approximately 13-fold with mix-hcDNA (**Fig. 2b**).

**Fig. 2.**
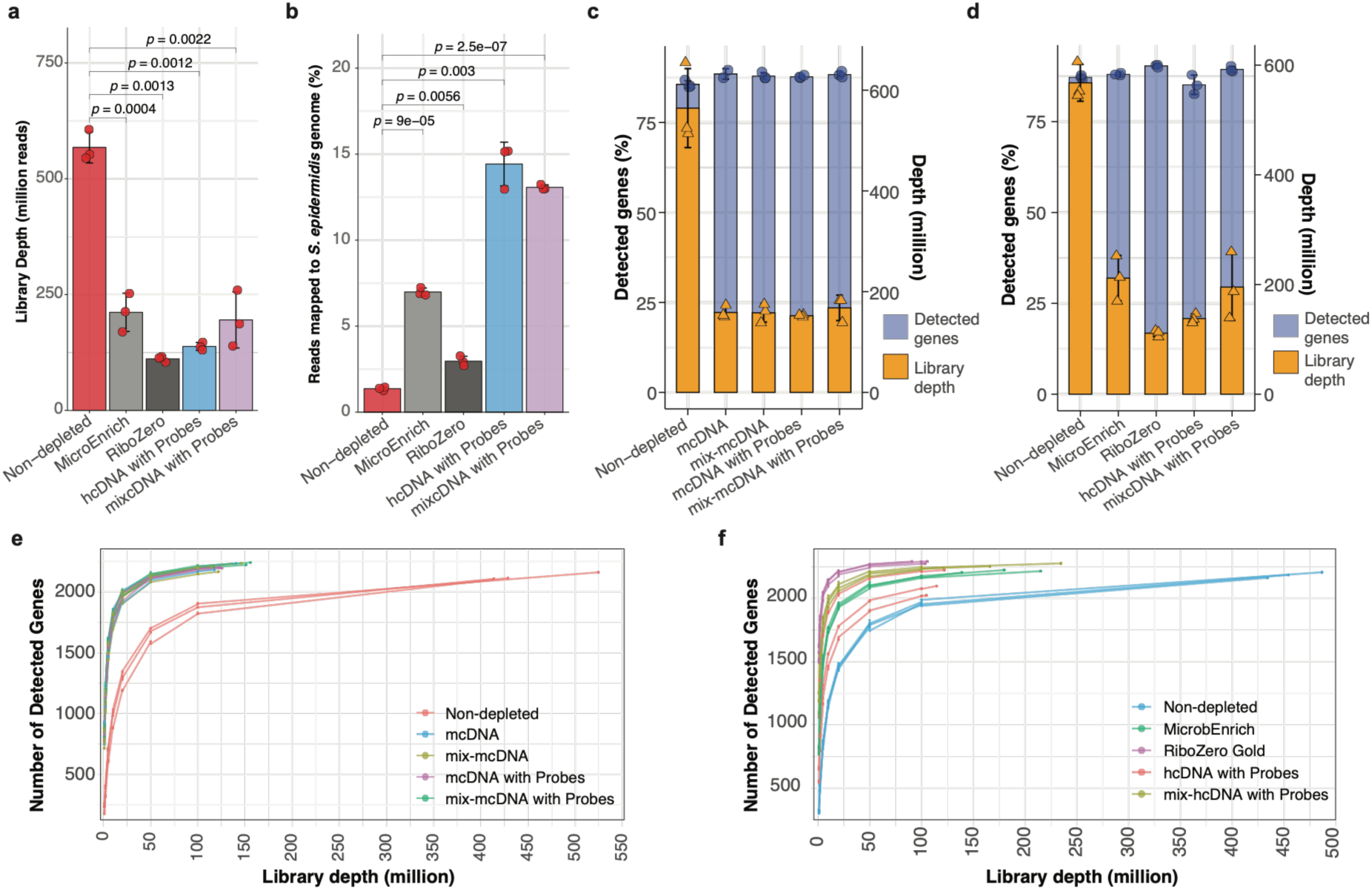
cDNA-guided depletion outperforms commercial enrichment strategies and reduces sequencing depth requirements for complete bacterial transcriptome coverage. **(a)** Library sequencing depth (million reads) for non-depleted, MICROBEnrich, RiboZero Gold, hcDNA with probes, and mix-hcDNA with probes conditions in human RNA mixtures. Data represent mean ± SEM; individual replicates shown as red dots. **(b)** Percentage of reads mapping to the *S. epidermidis* genome across human RNA mixture conditions. Data represent mean ± SEM; individual replicates shown as red dots. FDR-adjusted Welch’s t-test (**p < 0.01, ***p < 0.001). **(c, d)** Stacked bar plots showing the percentage of *S. epidermidis* genes detected (blue, left axis) and library sequencing depth (orange, right axis) at maximum sequencing depth for mouse **(c)** and human **(d)** mixtures across all depletion conditions. Individual replicates shown as triangles. **(e, f)** Rarefaction curves showing the number of detected *S. epidermidis* genes as a function of library depth for mouse **(e)** and human **(f)** mixtures. *S. epidermidis* contains 2,352 CDSs.

### Sequencing depth requirements for complete bacterial transcriptome coverage are markedly reduced by cDNA-guided host RNA depletion

Bacterial RNA recovery scaled proportionally with sequencing depth across all conditions, and the proportion of reads mapping to *S. epidermidis* per million total reads correlated strongly with the overall mapping percentage in both human and mouse datasets (**Supplementary Fig. S4**). We next quantified the fraction of *S. epidermidis* genes detected as a function of sequencing depth. In non-depleted samples, comprehensive gene detection required more than 600 million reads, with detection of rare transcripts expected to require billions (**Fig. 2c and d)**. With cDNA-guided depletion, the same level of detection was reached with ∼200 million reads in both mouse and human samples (**Fig. 2c and d**). Notably, in human samples RiboZero Gold achieved slightly higher gene detection at lower sequencing depth than the cDNA-guided methods. Rarefaction analysis confirmed that fully saturated bacterial transcriptome coverage was reached at ∼50–100 million reads in enriched samples for both mouse and human (**Fig. 2e and f**), representing a substantial reduction in sequencing cost and depth compared with non-depleted libraries.

### cDNA-guided host RNA depletion targets all major host RNA biotypes

To determine whether cDNA-guided depletion removes a broad spectrum of host RNAs beyond rRNA, we mapped reads from all conditions to the human or mouse genome and assigned them to annotated RNA biotypes, then compared the resulting compositions. In non-depleted mouse mixtures, libraries were dominated by rRNA, with additional contributions from non-coding classes including misc_RNA, snRNA, snoRNA, retained-intron transcripts, and NMD targets. After mcDNA or mix-mcDNA depletion, with or without probe supplementation, this composition was substantially redistributed, with protein-coding transcripts and rRNA becoming the most abundant biotypes (**Fig. 3a**). Human non-depleted mixtures showed a similar rRNA-dominated profile, which shifted toward a protein-coding-dominated composition after RiboZero treatment and after probe-supplemented hcDNA or mix-hcDNA depletion (**Fig. 3b**). Microb*Enrich*-treated samples were a partial exception: rRNA remained the most abundant biotype, although its proportion was reduced relative to protein-coding transcripts compared with the non-depleted baseline (**Fig. 3b**). Across both species, the relative proportions of every biotype shifted measurably in depleted versus non-depleted samples, consistent with broad-spectrum depletion rather than rRNA-specific removal (**Fig. 3a and b**).

**Fig. 3.**
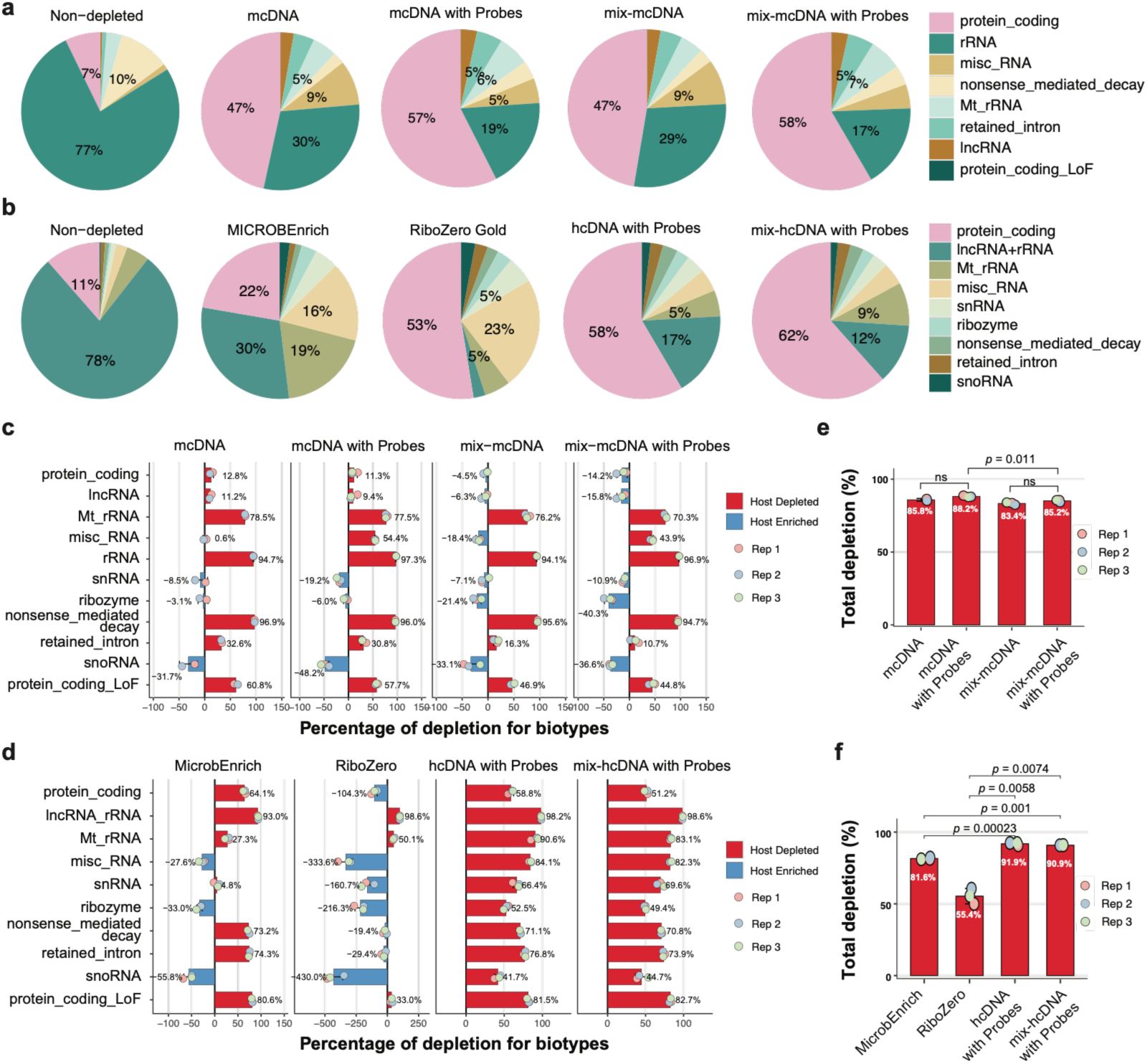
cDNA-guided host RNA depletion targets all major host RNA biotypes. **(a, b)** Pie charts showing the proportional composition of mapped reads by RNA biotype in mouse **(a)** and human **(b)** RNA mixtures across all depletion conditions. **(c, d)** Horizontal bar plots showing the percentage of depletion for each host RNA biotype, calculated using a pathogen-referenced normalization strategy, for mouse **(c)** and human **(d)** mixtures. Red bars indicate host-depleted biotypes and blue bars indicate relatively enriched biotypes. Individual biological replicates are shown as colored dots. **(e, f)** Total host RNA depletion efficiency (%) aggregated across all biotypes for mouse **(e)** and human **(f)** conditions. In mouse samples, total depletion ranged from 83.4% to 88.2%. In human samples, hcDNA and mix-hcDNA with probe supplementation achieved 90.9–91.9% total depletion. Data represent individual replicates; horizontal bars indicate group medians. FDR-adjusted Welch’s t-test; ns, not significant.

To evaluate depletion efficiency without the confounding effects of compositional shifts in highly depleted RNA-seq libraries, we used a pathogen-referenced normalization strategy where the absolute read counts mapped to the *S. epidermidis* genome were used as an internal anchor across all biological replicates as the depletion does not affect bacterial RNAs (**Supplementary Note S2, Supplementary Fig S5**). For each biological replicate, the host-to-pathogen ratio (*R*_ℎ*p*_) of a given biotype was calculated as:

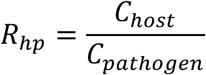

where *C*_ℎ*ost*_ is the read count mapped to a specific host biotype (rRNA, protein-coding mRNA, lncRNA, etc.) and *C_pat_*_ℎ*o*g*en*_ is the read count mapped to the *S. epidermidis* genome. The mean host-to-pathogen ratio of the non-depleted control samples (*R̄_baseline_*) was used as the reference. The percentage of depletion (*D*_%_) for each target replicate was computed as:

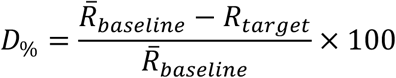

Positive *D*_%_ values indicate depletion of host RNAs relative to the pathogen background. Negative values denote enrichment of a biotype within the proportionally freed sequencing space. In mouse mixtures, cDNA-guided depletion efficiently removed the most abundant host RNA biotypes, both with and without probe supplementation, in mcDNA and mix-mcDNA conditions (**Fig. 3c**). Depletion of the misc_RNA biotype, which includes 7S RNA, was notably more efficient in probe-supplemented mcDNA and mix-mcDNA depletions, confirming the contribution of the 7S RNA-targeting probes. Protein-coding transcripts were not depleted by mix-mcDNA with or without probe supplementation, in contrast to the other conditions; the reason for this is unclear and requires further investigation (**Fig. 3c**). In human mixtures, hcDNA and mix-hcDNA with probe supplementation depleted all of the most abundant biotypes (**Fig. 3d**). RiboZero Gold, in contrast, depleted only rRNA, mitochondrial rRNA, and one minor biotype (protein_coding_LoF), while the remaining biotypes were relatively enriched (**Fig. 3d**). This apparent enrichment likely reflects RiboZero Gold’s known co-depletion of bacterial rRNA, which reduces the pathogen mapped reads and inflates the apparent host-to-pathogen ratio of all non-rRNA biotypes. MICROB*Enrich* depleted the majority of abundant host biotypes, but its overall efficiency was lower than that of the cDNA-guided methods, with the lowest rRNA depletion of any condition and a relative enrichment of misc_RNA (**Fig. 3d**). Together, these results show that cDNA-guided depletion, combined with supplemental probe targeting, depletes all major host RNA biotypes more comprehensively than the current commercial methods.

To assess depletion of overall host RNAs, reads from all targeted host biotypes within a given sample were aggregated into a total host pool (*C*_ℎ*ost*,*total*_):

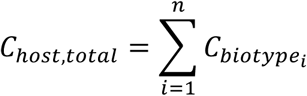

The total host-to-pathogen ratio was then calculated for each replicate as:

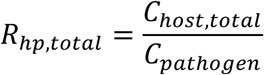

Using the mean aggregated ratio of the non-depleted baseline (*R̄_baseline_*_,*total*_), the total host depletion percentage was calculated as:

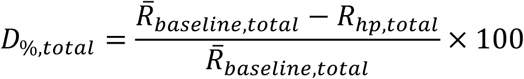

Total host RNA depletion ranged from 83.4% to 88.2% in mouse samples treated with mcDNA or mix-mcDNA (**Fig. 3e**). Although the differences between probe-supplemented mcDNA and mix-mcDNA were not statistically significant, mcDNA with probe supplementation achieved the highest efficiency in mouse samples (88.2%; Fig. 3e). In human samples, hcDNA and mix-hcDNA with probe supplementation achieved 90.9% to 91.9% total depletion (**Fig. 3f**). Both cDNA-guided approaches depleted host RNA significantly more efficiently than MICROB*Enrich* and RiboZero Gold, noting that part of the apparent gap for RiboZero Gold reflects its co-depletion of bacterial rRNA rather than poor host depletion *per se* (**Fig. 3f**). Together, these results show that cDNA-guided depletion efficiently removes the most abundant host RNA biotypes in both mouse and human backgrounds and outperforms two current commercial methods.

### cDNA-guided host RNA depletion preserves bacterial transcript coverage and expression composition

We asked whether our cDNA-guided host RNA depletion alters the representation of bacterial transcripts. We quantified expression level (TPM, Transcript Per Million) and evaluated *(i)* length-normalized coverage across the 5′ and 3’, stratified by CDS length (500–1,000 nt; 1,001–2,000 nt; 2,001–5,000 nt), and *(ii)* expression distributions across replicates and methods. Transcript coverage profiles closely overlapped for each length class, with no detectable 5′ or 3′ bias regardless of depletion or depletion methods in mouse or human samples (**Fig. 4a and b**). To test reproducibility and potential composition shifts, we performed pairwise comparisons among biological replicates within each treatment/sample. Pearson correlation coefficients exceeded 0.90 in all comparisons, indicating that cDNA-guided depletion maintains between-replicate similarity at least as well as non-depleted libraries (**Supplementary Fig. S6**). Finally, we examined whether depletion disproportionately impacts low- or high-abundance transcripts. Density plots of log₂(TPM+1) were highly similar between depleted and non-depleted samples. For mouse mixes, the expression distributions in mcDNA and mix-mcDNA samples, with or without supplemental rRNA probes, overlapped the non-depleted baseline almost completely. For human mixes, hcDNA and mix-hcDNA with probe supplementation produced similarly overlapping distributions (**Fig. 4c and d**). The same pattern was observed for MICROB*Enrich* and RiboZero Gold-treated samples, indicating that this preservation of transcript distribution is a general property of effective depletion rather than unique to our method. Together, these results show that cDNA-guided host RNA depletion enriches bacterial reads without distorting transcript coverage, length dependence, or abundance distributions, supporting quantitative downstream analyses such as differential expression in both human- and mouse-associated samples.

**Fig. 4.**
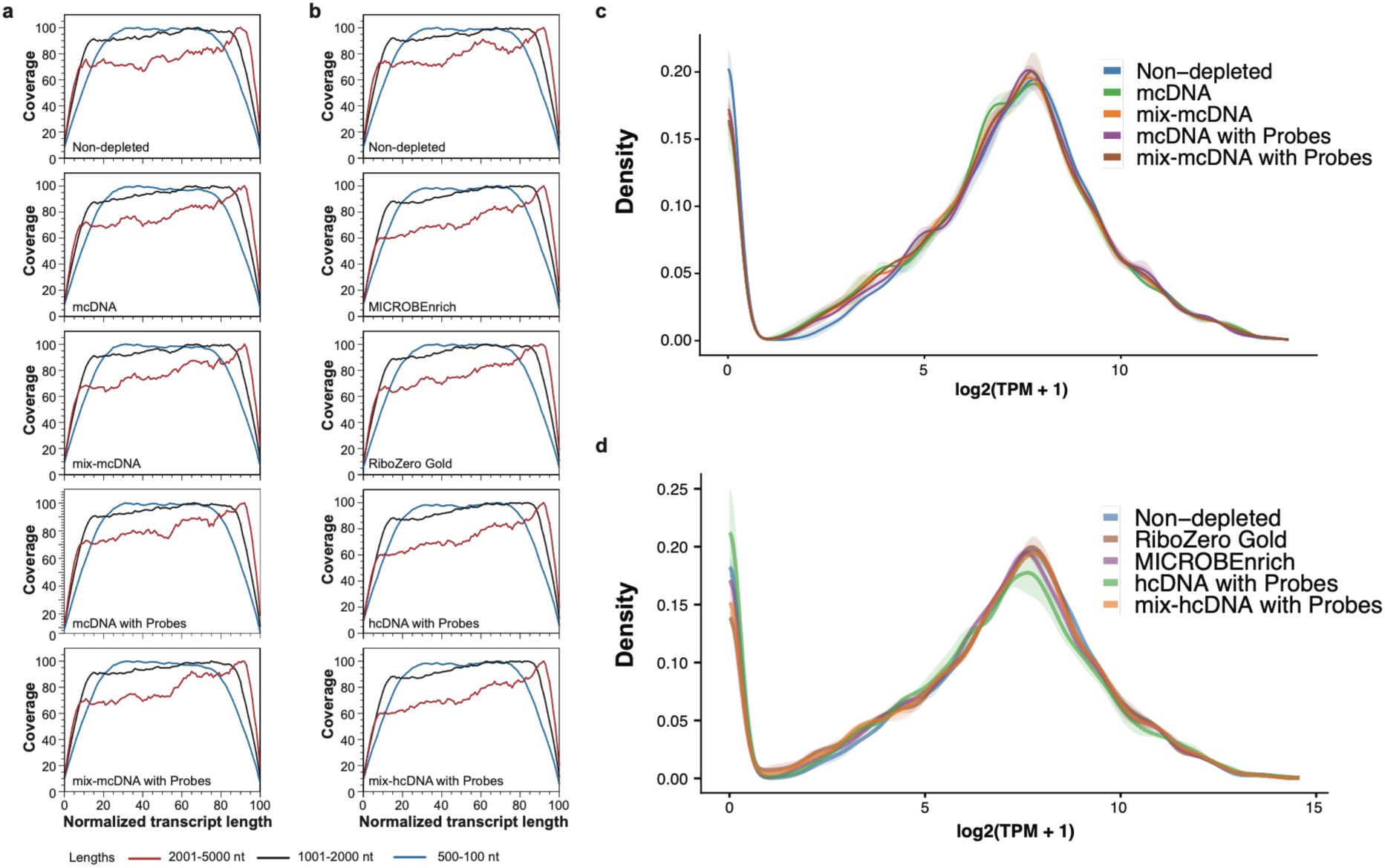
cDNA-guided host RNA depletion preserves bacterial transcript coverage and expression composition. **(a, b)** Length-normalized coverage profiles of *S. epidermidis* transcripts across the 5’→3’ axis, stratified by coding sequence (CDS) length (500–1,000 nt, blue; 1,001–2,000 nt, black; 2,001–5,000 nt, red), for mouse **(a)** and human **(b)** RNA mixtures across all depletion conditions. **(c, d)** Density plots of log₂(TPM+1) values showing the global expression distribution of *S. epidermidis* transcripts in mouse **(c)** and human **(d)** RNA mixtures across all depletion conditions. Shaded ribbons represent variability across biological replicates.

### cDNA-guided host RNA depletion preserves bacterial transcriptome integrity and differential responses

We next asked whether depleting host RNA alters the bacterial transcriptome itself or confounds downstream inference. To account for potential variations in sequencing depth for *S. epidermidis* across depleted and non-depleted host mixtures, bacterial RNA-only samples were included as baseline controls for more robust transcriptomic comparisons (**Supplementary Note S3**). Using log2-transformed TPM for all genes, we observed strong correlations between depleted libraries and both bacteria-only and non-depleted mixtures in human and mouse backgrounds, with replicate correlation comparable to (and often matching) non-depleted controls (**Fig. 5a and b**). To place these profiles in a broader transcriptional context, we embedded our datasets alongside 11 infection-relevant *S. epidermidis* transcriptomes from prior work^32^ using t-SNE. All depleted and non-depleted libraries clustered tightly with the corresponding bacteria-only controls and were clearly separated from stress-condition transcriptomes, indicating that depletion does not introduce an artificial global transcriptional signature (**Fig. 5c and d**). We then tested whether depletion affects biological interpretation under perturbation by performing differential expression for low-iron stress using limma–voom (TMM normalization, voom precision weights, empirical-Bayes moderation) and comparing each to the reference analysis (low iron versus bacteria-only). We defined DEGs as genes with FDR < 0.05 and |log2FC| > 1. In human mixtures, while cDNA-guided depletion with mix-hcDNA yielded the highest correlation in DEGs and directionality relative to the reference, exceeding RiboZero Gold and MICROB*Enrich*, cDNA-guided depletion with hcDNA showed comparable results (**Fig 5e**). In mouse mixtures, the mix-mcDNA condition supplemented with targeted probes produced the closest match, with other cDNA-guided conditions also outperforming comparator methods (**Fig. 5f**). Together, these analyses show that cDNA-guided host RNA depletion enriches bacterial reads without distorting global expression patterns or perturbation-responsive genes, supporting accurate downstream discovery at substantially reduced sequencing depth.

**Fig. 5.**
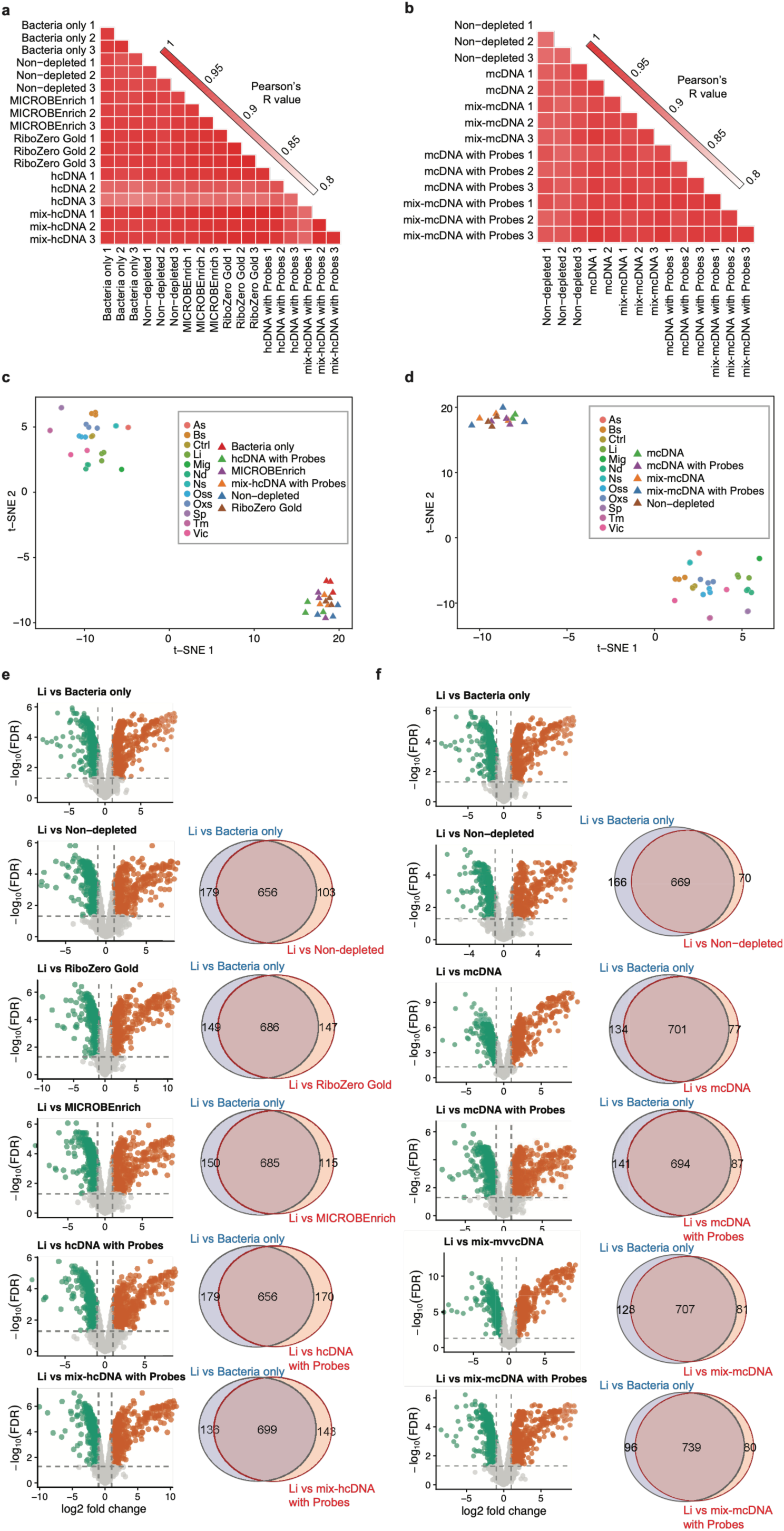
cDNA-guided host RNA depletion preserves global expression structure and differential expression fidelity. **(a, b)** Pearson correlation coefficients of bacterial transcriptomes between sample conditions and bacteria-only controls in human **(a)** and mouse **(b)** mixtures. **(c, d)** t-SNE embedding of all libraries from this study alongside 11 previously reported *S. epidermidis* transcriptomes measured under infection-relevant stress conditions in human **(c)** and mouse **(d)** mixtures. Stress conditions: acidic stress (As), bile stress (Bs), control (Ctrl), low iron (Li), microaerophilic growth (Mig), nutritional downshift (Nd), nitrosative stress (Ns), osmotic stress (Oss), oxidative stress (Oxs), stationary phase (Sp), temperature stress (Tm), and virulence-inducing condition (Vic). **(e)** Volcano plots of differential expression analysis under low iron (Li) versus bacteria-only, non-depleted, hcDNA, mix-hcDNA, RiboZero Gold, and MICROBEnrich conditions, using limma–voom. Points are colored by direction (upregulated, red; downregulated, blue) with FDR < 0.05 and |log₂ fold-change| > 1. **(f)** Volcano plots of differential expression analysis as in **(e)** using Li versus bacteria-only, non-depleted, mcDNA, mix-mcDNA, mcDNA with probes, and mix-mcDNA with probes. In **(e)** and **(f)**, dashed lines indicate significance thresholds and Venn diagrams show common and unique DEGs between comparisons.

### *In vivo* validation of cDNA-guided depletion using *Salmonella* infected mice

Having demonstrated the efficiency of our cDNA-guided depletion approach using defined bacterial–host RNA mixtures, we next sought to validate this method under physiologically relevant conditions. To this end, we applied our depletion strategy to *in vivo* samples derived from *Salmonella enterica* serovar Typhimurium-infected mice. Unlike the controlled spiking experiments described above, *in vivo* infection models present additional complexity, including variable bacterial loads, heterogeneous host immune responses, and a broader dynamic range of both host and bacterial transcripts. We employed two well-established mouse models of *S.* Typhimurium infection. In the first model, BALB/c mice, which are genetically susceptible to *Salmonella* infection due to a non-functional *Nramp1* allele, were pretreated with streptomycin 24 hours prior to infection to abolish microbiota-mediated colonization resistance, leading to acute colitis in the large intestine ^33,34^. Mice were then infected intragastrically with 5×10³ CFUs of *S.* Typhimurium, and cecal tissues were collected 24 hours post-infection. In the second model, CBA/Ca mice, which carry a functional *Nramp1* allele conferring *Salmonella* resistance, were used to model chronic infection. Following the same streptomycin pretreatment strategy, mice were infected with a higher dose of 5×10⁶ CFUs, cecal tissues were collected at day 2 and day 42 post-infection. These time points capture both the early and prolonged phases of bacterial colonization of cecum and shedding in feces which are characteristic of this model^35,36^.

For host RNA depletion, cDNA was generated from RNA isolated from uninfected mouse mesenteric lymph nodes (MLN) as it is a more sterile tissue free of microbiota in contrust and subsequently hybridized to RNA extracted from cecum samples collected from both infection models. Host RNA within the resulting RNA:cDNA hybrids were then selectively digested using RNase H, followed by DNase treatment to remove residual cDNA, and the resulting bacterial-enriched RNA was used for RNA-seq library preparation.

Given the complexity of *in vivo* samples and the expected variability in bacterial load across infection stages, non-depleted samples were sequenced to greater depths than in the spiked-in experiments, averaging 604 million, 566 million, and 1.8 billion reads for the 24-hour, day 2, and day 42 time points, respectively (**Fig. 6a**). Depleted samples required substantially fewer reads, with sequencing depths of 172 million, 107 million, and 810 million reads for the corresponding time points (**Fig. 6a**), reflecting the reduced sequencing burden conferred by host RNA removal.

**Fig. 6.**
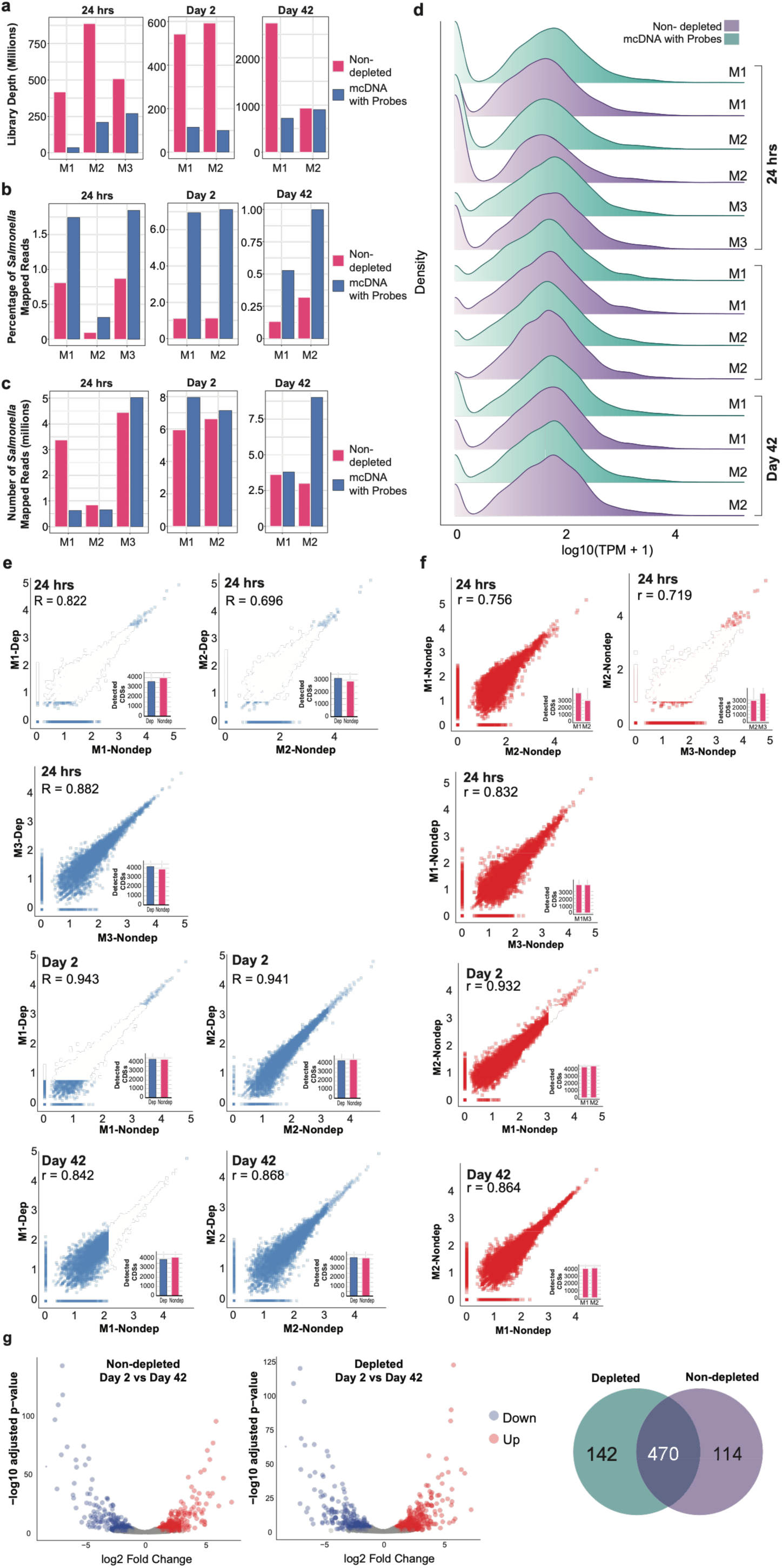
cDNA-guided host RNA depletion faithfully preserves the *S.* Typhimurium transcriptome across *in vivo* infection stages. **(a)** Library sequencing depth (millions of reads) for non-depleted (red) and mcDNA-depleted (blue) samples across biological replicates at 24 hours, day 2, and day 42 post-infection. **(b)** Percentage of reads mapping to the *S.* Typhimurium SL1344 genome across biological replicates for non-depleted and depleted conditions at each infection time point. **(c)** Absolute number of *S.* Typhimurium mapped reads (millions) per replicate for non-depleted and depleted conditions across infection time points. **(d)** Density plots of log₁₀(TPM+1) values for non-depleted (purple) and mcDNA-depleted (teal) samples across all replicates and time points at 24 hours, day 2, and day 42. **(e)** Pairwise scatterplots of log₁₀(TPM+1) values comparing depleted versus non-depleted samples within each biological replicate at 24 hours (R=0.822, R=0.696, R=0.882), day 2 (R=0.943, R=0.941), and day 42 (R=0.842, R=0.868). Inset bar plots show the number of detected CDSs in non-depleted and depleted conditions per replicate. **(f)** Pairwise scatterplots comparing non-depleted replicates against each other at 24 hours (r=0.756, r=0.719, r=0.832), day 2 (r=0.932), and day 42 (r=0.864). **(g)** Volcano plots of differential expression analysis comparing day 2 versus day 42 time points in non-depleted (left) and depleted (right) samples, showing upregulated (red) and downregulated (blue) genes with FDR < 0.05 and |log₂FC| > 1. Venn diagram shows 470 DEGs shared between conditions, with 142 and 114 DEGs detected exclusively in depleted and non-depleted samples, respectively.

In non-depleted samples, the proportion of reads mapping to the *S.* Typhimurium genome varied across infection stages, consistent with differences in bacterial burden: approximately 0.59% at 24 hours, 1.1% at day 2, and 0.22% at day 42 (**Fig. 6b**), corresponding to an average of 2.8, 6.2, and 3.3 million mapped reads (**Fig. 6c**). Following cDNA-guided depletion, bacterial read fractions increased to 1.3%, 7%, and 0.76% at the respective time points (**Fig. 6b**), with average mapped read counts of 2.1, 7.5, and 6.4 million reads (**Fig. 6c and Supplementary Dataset S1**), representing meaningful enrichment across all conditions. Notably, while the enrichment at 24 hours resulted in a comparable absolute mapped read count despite the increased read fraction, depletion at day 2 and day 42 yielded substantially higher absolute bacterial read numbers at markedly reduced sequencing depths, highlighting the practical efficiency gains of the approach. The more modest enrichment observed at 24 hours likely reflects the lower bacterial load characteristic of the early acute infection phase, which may limit the absolute gain achievable by depletion alone.

To further assess transcriptome-wide concordance between depleted and non-depleted samples, we examined the distribution of gene expression levels using density plots of log_10_(TPM+1) values. The distributions were highly similar between conditions across all time points (**Fig. 6d**), indicating that host RNA depletion does not introduce systematic bias in bacterial transcript quantification. Despite differences in sequencing depth and depletion efficiency across time points, gene coverage remained remarkably consistent between depleted and non-depleted samples. In non-depleted samples, 3,600, 4,387, and 4,136 *S.* Typhimurium CDSs were detected at 24 hours, day 2, and day 42, respectively, out of a total of 4,530 annotated CDSs in the SL1344 genome. Depleted samples yielded comparable coverage, with 3,656, 4,342, and 4,055 CDSs detected at the corresponding time points (**Fig. 6e and f**).

To further evaluate the concordance between depleted and non-depleted samples at the individual replicate level, pairwise correlation analysis was performed on TPM values across all time points. At day 2, correlation coefficients were consistently high (r=0.943 and r=0.941), indicating strong agreement between depleted and non-depleted transcriptional profiles (**Fig. 6e**). Similarly, day 42 samples showed robust correlations (r=0.842 and r=0.868), demonstrating that depletion faithfully preserves the transcriptional landscape even during chronic infection. At 24 hours, correlations were more variable across replicates (r=0.822, r=0.696, and r=0.882), which is consistent with the density plot distributions showing pronounced zero inflation at this time point (**Fig. 6e**). This zero inflation likely reflects the lower bacterial load during early acute infection, where a substantial proportion of genes are expressed at very low or undetectable levels, increasing sensitivity to technical variability between replicates. To contextualize these values, pairwise correlations between non-depleted replicates within the same time point ranged from r=0.719 to r=0.832 at 24 hours, r=0.932 at day 2, and r=0.864 at day 42 (**Fig. 6f**), indicating that the concordance between depleted and non-depleted samples is comparable to or exceeds the natural between-replicate variability observed within non-depleted libraries alone. Notably, the lower replicate number at day 2 and day 42 (n=2) due to limited sample availability may constrain the statistical power at these time points, though the high correlation values observed suggest reliable concordance between conditions.

To evaluate whether depletion influences the detection of differentially expressed genes (DEGs), we compared transcriptional profiles between day 2 and day 42 in both depleted and non-depleted samples. This analysis identified 470 DEGs shared between the two conditions, demonstrating broad concordance in the biological signal captured by both approaches. However, 142 DEGs were detected exclusively in depleted samples and 114 exclusively in non-depleted samples (**Fig. 6g**), suggesting that host RNA depletion improves sensitivity for detecting differentially expressed bacterial transcripts, particularly those of lower abundance that may be obscured by residual host RNA in non-depleted libraries. Collectively, these findings demonstrate that cDNA-guided host RNA depletion preserves comprehensive bacterial transcriptome coverage, maintains transcriptome-wide expression distributions, and enhances the detection of biologically relevant differential gene expression, while substantially reducing the sequencing depth required, confirming the robustness of the approach across distinct infection models and disease stages.

## Discussion

Capturing the pathogen transcriptome during active infection is bottlenecked by the overwhelming dominance of host RNA. While standard commercial kits successfully remove ribosomal RNA (rRNA), they leave an expansive pool of host messenger and non-coding RNAs that outcompete rare bacterial transcripts for sequencing reads.

A review of recent *in vivo* transcriptomic literature underscores the severity of this limitation. Our previous work on *Yersinia pseudotuberculosis* infection in mouse model yielded as little as 0.0016% Yersinia transcripts despite enrichment with MICROBEnrich and MICROBExpress^37^. Further study with utilizing anti-*Yersinia* antibody-coupled beads to isolate intact bacterial cells, we achieved a 43-fold increase in bacterial transcript recovery (0.07%). Despite this improvement, the high host-to-pathogen RNA ratio necessitated an ultra-deep sequencing depth of 600 million reads per library to ensure sufficient coverage for differential expression analysis^31^. In a model of *Leptospira interrogans* infection, standard rRNA depletion yielded a bacterial transcript fraction of only 4.57% at 40 million reads, necessitating deep sequencing (400 million reads) to achieve sufficient genome coverage^38^. Similarly, in clinical *Pseudomonas aeruginosa* infections, overwhelming host background in chronic wounds suppressed bacterial read fractions to as low as 0.0048%, necessitating the bioinformatic exclusion of thousands of genes to salvage the analysis^39^. Even advanced dual RNA-seq studies of *Streptococcus pneumoniae* in murine lungs yielded bacterial read fractions of only 0.06% to 1.7% in different samples^40^, while studies of *Escherichia coli* (ETEC) and *Staphylococcus aureus* similarly struggled with low abundance recovery despite rigorous rRNA depletion^41,42^. Furthermore, early clearance of uropathogenic *E. coli* within macrophages rendered transcriptomic analysis completely impossible due to sequencing drop-out^43^.

To overcome this universal technical barrier, we developed an adaptive, cDNA-guided broad host RNA depletion method that selectively eliminates host-derived transcripts directly from total host-microbe RNA mixtures. By utilizing cDNAs derived from the sample itself to guide RNase H digestion, this approach adapts to the sample’s specific host transcriptome without the need for custom probe design. Such efficient target RNA enrichment fundamentally redefines the economic and computational feasibility of *in vivo* transcriptomics. In non-depleted samples, complete detection of the bacterial genome required over 600 million reads. With cDNA-guided depletion, fully saturated bacterial transcriptome coverage is achieved at merely 50 to 100 million reads. Using cDNA generated from infected sample RNA, host transcripts are selectively depleted while bacterial RNA remains intact. Because host transcripts vastly outnumber bacterial ones, reverse transcriptase becomes fully saturated during cDNA synthesis, making bacterial cDNA generation highly unlikely. Subsequently, RNase H digestion efficiently degrades the abundant host RNA:cDNA hybrids, while the rare bacterial hybrids remain largely uncleaved within the short digestion window. The bacterial transcriptome is therefore preserved and enriched for downstream analysis.

Unlike standard commercial enrichment methods that actively co-deplete bacterial ribosomal RNA, our cDNA-guided strategy specifically targets the host transcriptome, preserving the bacterial rRNA pool and its biological information. The quantity and quality of bacterial rRNA directly reflect the pathogen’s metabolic state and *in vivo* growth rate, as ribosome concentrations correlate strongly with ongoing microbial replication^44^. Consequently, tracking steady-state rRNA levels serves as both a reliable proxy for physiological adaptation and an essential biomarker to distinguish viable, metabolically active bacteria from dead cells or nucleic acid debris within infected tissues^45^. This preservation is particularly crucial during infection, where severe host-induced stresses frequently force pathogens into viable but non-culturable or persister phenotypes characterized by drastically altered resource allocation and translation dynamics^46^. By comprehensively sequencing the entire intact bacterial RNA pool, our methodology transforms a perceived technical byproduct into a highly valuable dataset for evaluating pathogen survival, growth kinetics, and physiological vitality directly at the infection site.

If applied to previous *in vivo* studies, this methodology would have transformed data yields. For example, the *L. interrogans* study^38^ could have achieved near-complete genome coverage using one-eighth of their sequencing effort. In practical terms, it reduces sequencing cost by 88%-98% per sample. Furthermore, physically reducing the uninformative read count by up to 95% circumvents immense computational bottlenecks, cutting read-alignment times from hours down to minutes and drastically reducing cloud storage demands.

The sample-adaptive nature of cDNA-guided depletion offers usage beyond human bacterial pathogenesis. In veterinary and agricultural science, dual RNA-seq is frequently paralyzed by poorly annotated host reference genomes, causing massive in silico cross-mapping errors. Because our RNase H-mediated depletion relies on physical sequence homology within the sample rather than computational databases, it effortlessly bypasses the need for perfectly annotated host genomes.

Similarly, this methodology is theoretically able to transform viral transcriptomics. During acute viral infections or clinical sepsis diagnostics, rare viral RNA or pathogenic transcripts are dominated by host mRNA. Because viral genomes are highly divergent from eukaryotic hosts, the host-derived cDNA probes could exclusively target and erase the host background without hybridizing to the viral RNA. This targeted depletion enables the sequencing of ultra-low titer viruses directly from clinical samples, facilitating unbiased detection of emerging infectious diseases and high-resolution mapping of viral transcriptomes.

Ultimately, our sample-derived cDNA depletion establishes a culture-independent, cost-aware standard that shifts the limiting factor in infection biology from the host background to the biological signal itself.

## Material and Methods

### Animal Ethics and Housing

Mice were housed in the infection unit at the Umeå Centre for Comparative Biology (UCCB) in accordance with the Swedish National Board for Laboratory Animals guidelines. All animal procedures were approved by the Animal Ethics Committee of Umeå University (Dnr A22/2021). Upon arrival, mice were acclimated for one week prior to experiments. Mice were maintained on a 12-hour light/dark cycle and fed food and water *ad libitum*.

### Mouse Infection Models

BALB/c and CBA/Ca mice (8-9 weeks old) were pretreated with streptomycin (20 mg in 0.1 mL sterile PBS, intragastrically) 24 hours prior to infection. *S.* Typhimurium SL1344 was grown overnight, diluted 1:40, and incubated for 4 hours to reach late exponential phase, then diluted in PBS to the desired inoculum concentration. BALB/c mice were infected intragastrically with 5×10³ CFUs in 0.1 mL PBS, and cecal tissues were collected 24 hours post-infection after cecal content were removed by washing cecal tissues several times in RNA protect. CBA/Ca mice were infected intragastrically with 5×10⁶ CFUs in 0.1 mL PBS, and cecal tissues were collected at day 2 and day 42 post-infection.

### RNA extraction from Bacteria

Total RNA from *S. epidermidis* 1457 was extracted using TRIzol™ reagent (Invitrogen, Life Technologies, CA, USA) following the manufacturer’s instructions. Bacteria were cultured in tryptic soy broth (TSB) at 37 °C until mid-logarithmic phase. A 5 mL aliquot of the culture was harvested by centrifugation at maximum speed in a benchtop centrifuge for 3 min. The resulting pellet was subjected to TRIzol extraction, and the upper aqueous phase was further processed using the Direct-zol™ RNA Miniprep Kit (Zymo Research, USA) according to the manufacturer’s protocol. To eliminate contaminating genomic DNA, the purified RNA was treated with TURBO™ DNase (Invitrogen, Life Technologies, CA, USA) as per the manufacturer’s instructions. DNase-treated RNA was subsequently cleaned and concentrated using the RNA Clean & Concentrator™ Kit (Zymo Research). RNA yield and purity were determined spectrophotometrically using a NanoDrop instrument (Thermo Fisher Scientific), and RNA integrity was assessed by Fragment analyser system (Agilent).

### RNA Extraction from Mammalian Cell Lines

Caco-2 cells (ATCC) were maintained in RPMI-1640 medium (Sigma-Aldrich, USA) supplemented with 10% (v/v) fetal bovine serum (FBS), 1% (v/v) penicillin–streptomycin, and non-essential amino acids. Cells were cultured at 37 °C in a humidified atmosphere containing 5% CO₂ and subcultured at 80–90% confluence every 3–4 days. For RNA extraction, near-confluent cells were detached with 0.25% trypsin–EDTA, collected by centrifugation at 120 × *g* for 10 min at room temperature, and immediately processed using TRIzol reagent as described above.

### RNA Extraction from Tissues

Mesenteric lymph node (MLN) tissues were collected and immediately snap-frozen in liquid nitrogen. Frozen tissues were homogenized in TRIzol reagent using a mechanical tissue homogenizer. RNA was isolated through phase separation according to the procedure described for bacterial samples.

### cDNA Synthesis

Complementary DNA (cDNA) was synthesized from total host RNA (isolated from cell cultures or tissues) or mixed RNA samples (host + bacterial RNA) using the SuperScript™ IV First-Strand Synthesis System (Thermo Fisher Scientific), selected based on a comparative efficiency assessment of five commercially available reverse transcriptases in which it yielded the highest cDNA output following RNase H digestion of the RNA template **(Supplementary Note S1, Supplementary Fig. S1).** Briefly, 3 µg of RNA was combined with 1 µL of 0.2 µg µL⁻¹ random hexamers and 1 µL of 10 mM dNTP mix in a total volume of 14 µL. The mixture was incubated at 65 °C for 5 min, followed by rapid cooling to 4 °C for 1 min. Subsequently, 4 µL of 5X first-strand buffer, 1 µL of 0.1 M DTT, and 1 µL of SuperScript IV reverse transcriptase (200 U µL⁻¹) were added to bring the reaction volume to 20 µL. Reverse transcription was carried out at 50 °C for 30 min, followed by enzyme inactivation at 80 °C for 10 min.

Following cDNA synthesis, residual RNA templates were removed to prevent interference with downstream applications. To achieve this, 2 µL RNase H (5 U µL⁻¹), 2 µL RNase A (10 mg mL⁻¹), and 2.4 µL RNase H reaction buffer (10×) were added, and the reaction was incubated at 37 °C for 30 min before cooling to 4 °C. cDNA was subsequently purified using AMPure XP beads at 2× bead concentration. Purified cDNA was stored at −80 °C until further use (**Supplementary Note S4, Supplementary Fig. S7**).

### Enrichment of Bacterial RNA

For enrichment experiments host RNA was mixed with 1% bacterial RNA. Reaction mixture consisted of 400 ng of RNA (396 ng host RNA+4 ng bacterial RNA), 1 µg cDNA (obtained from either host or mixed RNA), 1x hybridization buffer (50 mM Tris-HCl pH 7.5, 100 mM NaCl) and 50 μM EDTA in a final volume of 20 μL. This mixture was then subjected to a controlled hybridization protocol and reaction performed at 55°C. Total 10μL Preheated RNase H mixture (1 µL Thermostable RNase H (5U/µL, NEB M0523S), 2 µL 5x hybridization buffer, 6 μL 50 mM MgCl_2_, and 1 μL nuclease-free water) and then added to the sample mix and incubated at 55°C for 10 minutes. Additional probes targeting the 7S RNA and 18S rRNA sequences were designed and added (50 nM final concentration each) to depletion mixture (**Supplementary Table S1**). The RNase H treated sample was subsequently cleaned and eluted in 20 µL nuclease-free water using the RNA Clean & Concentrator™ Kit (Zymo Research). cDNAs in the mixture were then degraded by adding 2 μL of TURBO^TM^ DNase (2U/μL, Invitrogen AM2238) and 2.4 μL of the supplied 10X Reaction Buffer to tubes, mixed, incubated at 37°C for 30 min, and placed on ice. Subsequently, the reaction mixture was subjected to column-based purification to isolate and clean up the enriched bacterial RNA. The concentrations of purified RNA samples were measured using the Qubit^™^ 4.0 fluorometer. RNA integrity before and after depletion was assessed by Bioanalyzer High Sensitivity RNA analysis (**Supplementary Fig. S8**)

### Enrichment of Bacterial RNA using commercially available kits

Using 1 µg of RNA (containing 1% bacterial RNA) Ribo-Zero Gold (Illumina, RS-122-2303) and MICROBEnrich (Ambion, AM1901) kits were used according to the manufacturer’s protocol to check efficiency of our method.

### RNA-seq library preparation

The enriched RNA samples were taken for library preparation using NEBNext Ultra™ II Directional RNA Library Prep kit according to the manufacturer’s protocol. Briefly, priming and fragmentation was performed by incubating RNA samples 94 °C for 15 min (undepleted samples) or 7 min (depleted samples) with random primers then transferring the tube to ice. First and second-strand cDNA synthesis, end-prep, and adaptor ligation were performed according to the manufacturer’s instructions. Finally, we constructed the libraries using the NEBNext Multiplex Oligos for Illumina kit, which is required for amplification of the ligated cDNA according to the manufacturer’s instruction. Indexed cDNA was purified 0.9× AMPure XP magnetic beads (Beckman Coulter). Libraries were quantified on a Qubit^™^ 4.0 fluorometer, and final library size was assessed by running libraries on Bioanalyzer High Sensitivity Chip (Agilent, 5067–4626) (**Supplementary Fig. S2**).

### In vivo RNA-seq data processing

Raw paired-end sequencing reads were preprocessed using Cutadapt^47^. The Illumina adapter sequence AGATCGGAAGAGC was removed from both mates using -a and -A. Bases below a Phred quality score of 20 were trimmed from the 5′ and 3′ ends of both reads (-q 20,20), terminal ambiguous nucleotides were removed (--trim-n), and reads shorter than 30 bp after trimming were discarded (-m 30).

Processed reads were initially aligned to the *Mus musculus* GRCm38 reference genome using STAR v2.7.1a^48^. The genome index was generated from chromosomes 1–19, X, Y, and MT using the Ensembl release 101 annotation (*Mus*_*musculus*.GRCm38.101.gtf) ^49^. Reads were aligned in paired-end mode using two-pass mapping (--twopassMode Basic). A maximum of 10 alignment loci per fragment was permitted (--outFilterMultimapNmax 10), and no more than five mismatches per paired-end fragment were allowed (--outFilterMismatchNmax 5). Alignments were written as coordinate-sorted BAM files, while reads classified as unmapped by STAR were exported as paired FASTQ files for subsequent bacterial alignment. The number of mouse-mapped reads was calculated as the sum of uniquely mapped reads and reads mapped to multiple loci. Reads mapping to more than 10 loci were classified by STAR as unmapped and were not included in the mouse-mapped total.

Host-unmapped reads were subsequently aligned to the *Salmonella enterica* subsp. *enterica* serovar Typhimurium SL1344 genome assembly ASM21085v2 (GCA_000210855.2) using Bowtie2 ^50^. Bowtie2 was run in paired-end, very-sensitive mode with mixed and discordant alignment modes disabled (--very-sensitive --no-mixed --no-discordant). Alignments were converted from SAM to BAM format and coordinate-sorted using SAMtools^51^.

Gene-level quantification was performed separately for mouse and *Salmonella* using featureCounts from Subread v2.0.3 ^52^ and organism-specific GTF annotation files. Paired-end fragments were counted using reverse-stranded counting specified by -s2. Multimapping alignments and alignments overlapping more than one gene were excluded because the -M and -O options were not enabled. Mouse reads were quantified using *Mus*_*musculus*.GRCm38.101.gtf, whereas *Salmonella* reads were quantified using the corresponding SL1344 ASM21085v2 annotation (sl1344.ASM21085v2.48.gtf) from Ensembl Bacteria release 48^53^.

## Data availability

The raw RNA-seq reads are available at NCBI GEO database with accession number: GSE331001.

## Code availability

No custom code was developed for this study. All analyses were performed using published software packages with default parameters unless otherwise specified, as described in the Materials and Methods section.

## Acknowledgements

This work was supported by Swedish Research Council (No. 2021-02466), Kempestiftelserna (JCK22-0017), and the Medical Faculty at Umeå University (FS 2.1.6-281-22) to K. Avican, by Swedish Research Council Excellence Center grant (No. 2022-06543) for the Center for Modeling Adaptive Mechanisms in Living Systems Under Stress to K. Avican. We acknowledge µNordic Single Cell Hub (µNiSCH), Genomic Medicine Center Norr (GMC Norr) and National Genomics Infrastructure in Stockholm funded by Science for Life Laboratory, the Knut and Alice Wallenberg Foundation and the Swedish Research Council. The authors acknowledge the computational and storage resources provided at PDC, HPC2N, and UPPMAX. Access to Dardel at PDC was provided through NAISS 2025/22-1408, as a continuation of NAISS 2024/22-1434, and through NAISS 2025/22-234, awarded to Leyden Fernandez Vidal as a continuation of NAISS 2023/22-977. Project NAISS 2023/22-977 also provided access to Rackham at UPPMAX and was itself a continuation of SNIC 2022/22-849, awarded to Barbara Forró. Access to Kebnekaise at HPC2N was provided through hpc2n2025-200, as a continuation of hpc2n2025-084. Storage at PDC was supported through NAISS 2025/6-387, as a continuation of NAISS 2024/6-335, and NAISS 2024/23-518, awarded to Leyden Fernandez Vidal as a continuation of NAISS 2023/23-485.

## Authors contributions

TD, OS, UA, RFF and KA conceived, designed, and performed the experiments and TD, OS, LF and KA developed the pipeline and analysed the data. TD, UA and KA wrote the manuscript with input from OS and LF.

## References

1 He, S. in Metagenomics for Microbiology 109–142 (Elsevier, 2025).

2 Wang, Z., Gerstein, M. & Snyder, M. RNA-Seq: a revolutionary tool for transcriptomics. Nat Rev Genet 10, 57–63 (2009). 10.1038/nrg2484

3 Giannoukos, G. et al. Efficient and robust RNA-seq process for cultured bacteria and complex community transcriptomes. Genome Biology 13, r23 (2012). 10.1186/gb-2012-13-3-r23

4 Pitashny, M. et al. NGS in the clinical microbiology settings. Frontiers in Cellular and Infection Microbiology Volume 12 - 2022 (2022). 10.3389/fcimb.2022.955481

5 Gioitta Iachino, S., Scaggiante, F., Mazzarisi, C. & Schaller, C. The Role of Next-Generation Sequencing (NGS) in the Relationship between the Intestinal Microbiome and Periprosthetic Joint Infections: A Perspective. Antibiotics 13, 931 (2024).

6 Westermann, A. J., Barquist, L. & Vogel, J. Resolving host–pathogen interactions by dual RNA-seq. Plos Pathog 13, e1006033 (2017). 10.1371/journal.ppat.1006033

7 Chen, Q. et al. Clinical diagnostic value of targeted next-generation sequencing for infectious diseases (Review). Mol Med Rep 30, 153 (2024). 10.3892/mmr.2024.13277

8 Ojala, T., Häkkinen, A.-E., Kankuri, E. & Kankainen, M. Current concepts, advances, and challenges in deciphering the human microbiota with metatranscriptomics. Trends in Genetics 39, 686–702 (2023). 10.1016/j.tig.2023.05.004

9 Prezza, G. et al. Improved bacterial RNA-seq by Cas9-based depletion of ribosomal RNA reads. Rna 26, 1069–1078 (2020).

10 Herbert, Z. T. et al. Cross-site comparison of ribosomal depletion kits for Illumina RNAseq library construction. BMC Genomics 19, 199 (2018). 10.1186/s12864-018-4585-1

11 Haas, B. J., Chin, M., Nusbaum, C., Birren, B. W. & Livny, J. How deep is deep enough for RNA-Seq profiling of bacterial transcriptomes? BMC Genomics 13, 734 (2012). 10.1186/1471-2164-13-734

12 Culviner, P. H., Guegler, C. K. & Laub, M. T. A Simple, Cost-Effective, and Robust Method for rRNA Depletion in RNA-Sequencing Studies. Mbio 11, 10.1128/mbio.00010-00020 (2020). https://doi.org:doi:10.1128/mbio.00010-20

13 Everaert, C. et al. Blocking Abundant RNA Transcripts by High-Affinity Oligonucleotides during Transcriptome Library Preparation. Biol Proced Online 25, 7 (2023). 10.1186/s12575-023-00193-3

14 Petrova, O. E., Garcia-Alcalde, F., Zampaloni, C. & Sauer, K. Comparative evaluation of rRNA depletion procedures for the improved analysis of bacterial biofilm and mixed pathogen culture transcriptomes. Scientific Reports 7, 41114 (2017). 10.1038/srep41114

15 Pang, X. et al. Bacterial mRNA purification by magnetic capture-hybridization method. Microbiol Immunol 48, 91–96 (2004). 10.1111/j.1348-0421.2004.tb03493.x

16 Chen, Z. & Duan, X. Ribosomal RNA depletion for massively parallel bacterial RNA-sequencing applications. Methods Mol Biol 733, 93–103 (2011). 10.1007/978-1-61779-089-8_7

17 Morlan, J. D., Qu, K. & Sinicropi, D. V. Selective depletion of rRNA enables whole transcriptome profiling of archival fixed tissue. PLoS One 7, e42882 (2012). 10.1371/journal.pone.0042882

18 Andrzejewska-Romanowska, A., Tykwińska, E., Śledziński, P. & Pachulska-Wieczorek, K. A comparative analysis of mRNA enrichment strategies and guidance for improving their efficiency. Scientific Reports 15, 17890 (2025). 10.1038/s41598-025-02082-z

19 Wahl, A., Huptas, C. & Neuhaus, K. Comparison of rRNA depletion methods for efficient bacterial mRNA sequencing. Scientific Reports 12, 5765 (2022). 10.1038/s41598-022-09710-y

20 He, S. et al. Validation of two ribosomal RNA removal methods for microbial metatranscriptomics. Nature Methods 7, 807–812 (2010). 10.1038/nmeth.1507

21 Betin, V. et al. Hybridization-based capture of pathogen mRNA enables paired host-pathogen transcriptional analysis. Scientific Reports 9, 19244 (2019). 10.1038/s41598-019-55633-6

22 Armour, C. D. et al. Digital transcriptome profiling using selective hexamer priming for cDNA synthesis. Nature Methods 6, 647–649 (2009). 10.1038/nmeth.1360

23 Huang, Y., Sheth, R. U., Kaufman, A. & Wang, H. H. Scalable and cost-effective ribonuclease-based rRNA depletion for transcriptomics. Nucleic Acids Research 48, e20–e20 (2019). 10.1093/nar/gkz1169

24 Wangsanuwat, C., Heom, K. A., Liu, E., O’Malley, M. A. & Dey, S. S. Efficient and cost-effective bacterial mRNA sequencing from low input samples through ribosomal RNA depletion. BMC genomics 21, 717 (2020).

25 Choe, D. et al. RiboRid: A low cost, advanced, and ultra-efficient method to remove ribosomal RNA for bacterial transcriptomics. PLOS Genetics 17, e1009821 (2021). 10.1371/journal.pgen.1009821

26 Qasim, M. S. & Sarin, L. P. An efficient one-step rRNA depletion method for RNA sequencing in non-model organisms. bioRxiv, 2025.2008.2015.670489 (2025). 10.1101/2025.08.15.670489

27 Marsh, J. W., Humphrys, M. S. & Myers, G. S. A. A Laboratory Methodology for Dual RNA-Sequencing of Bacteria and their Host Cells In Vitro. Front Microbiol 8, 1830 (2017). 10.3389/fmicb.2017.01830

28 Westermann, A. J., Gorski, S. A. & Vogel, J. Dual RNA-seq of pathogen and host. Nature Reviews Microbiology 10, 618–630 (2012). 10.1038/nrmicro2852

29 Robbe-Saule, M., Babonneau, J., Sismeiro, O., Marsollier, L. & Marion, E. An Optimized Method for Extracting Bacterial RNA from Mouse Skin Tissue Colonized by Mycobacterium ulcerans. Front Microbiol 8, 512 (2017). 10.3389/fmicb.2017.00512

30 Pisu, D., Huang, L., Grenier, J. K. & Russell, D. G. Dual RNA-Seq of Mtb-Infected Macrophages In Vivo Reveals Ontologically Distinct Host-Pathogen Interactions. Cell Rep 30, 335–350.e334 (2020). 10.1016/j.celrep.2019.12.033

31 Sarigöz, O. et al. 6S RNA facilitates bacterial virulence and adaptation at the epithelial barrier. bioRxiv, 2025.2010.2007.681022 (2025). 10.1101/2025.10.07.681022

32 Avican, K. et al. RNA atlas of human bacterial pathogens uncovers stress dynamics linked to infection. Nat Commun 12, 3282 (2021). 10.1038/s41467-021-23588-w

33 Barthel, M. et al. Pretreatment of mice with streptomycin provides a Salmonella enterica serovar Typhimurium colitis model that allows analysis of both pathogen and host. Infection and immunity 71, 2839–2858 (2003).

34 Walker, G. T., Gerner, R. R., Nuccio, S. P. & Raffatellu, M. Murine models of Salmonella infection. Current protocols 3, e824 (2023).

35 Grote, A. et al. Persistent Salmonella infections in humans are associated with mutations in the BarA/SirA regulatory pathway. Cell host & microbe 32, 79–92. e77 (2024).

36 Rogers, A. W. et al. Salmonella re-engineers the intestinal environment to break colonization resistance in the presence of a compositionally intact microbiota. Cell host & microbe 32, 1774–1786. e1779 (2024).

37 Avican, K. et al. Reprogramming of Yersinia from virulent to persistent mode revealed by complex in vivo RNA-seq analysis. Plos Pathog 11, e1004600 (2015). 10.1371/journal.ppat.1004600

38 Giraud-Gatineau, A., Haustant, G., Monot, M., Picardeau, M. & Benaroudj, N. In vivo dual RNA-Seq uncovers key effectors of epithelial barrier disruption by an extracellular pathogen. Nat Commun 17 (2026). 10.1038/s41467-026-69033-8

39 Cornforth, D. M. et al. Pseudomonas aeruginosa transcriptome during human infection. Proc Natl Acad Sci U S A 115, E5125–e5134 (2018). 10.1073/pnas.1717525115

40 Oh, M. W. et al. Time-resolved RNA-seq analysis to unravel the in vivo competence induction by Streptococcus pneumoniae during pneumonia-derived sepsis. Microbiol Spectr 12, e0305023 (2024). 10.1128/spectrum.03050-23

41 Crofts, A. A. et al. Enterotoxigenic E. coli virulence gene regulation in human infections. Proc Natl Acad Sci U S A 115, E8968–e8976 (2018). 10.1073/pnas.1808982115

42 Jin, Q. et al. Dual RNA-seq reveals the complement protein C3-mediated host-pathogen interaction in the brain abscess caused by Staphylococcus aureus. mSystems 10, e0154024 (2025). 10.1128/msystems.01540-24

43 Mavromatis, C. H. et al. The co-transcriptome of uropathogenic Escherichia coli-infected mouse macrophages reveals new insights into host-pathogen interactions. Cell Microbiol 17, 730–746 (2015). 10.1111/cmi.12397

44 Scott, M., Gunderson, C. W., Mateescu, E. M., Zhang, Z. & Hwa, T. Interdependence of cell growth and gene expression: origins and consequences. Science 330, 1099–1102 (2010). 10.1126/science.1192588

45 Cangelosi, G. A. & Meschke, J. S. Dead or alive: molecular assessment of microbial viability. Appl Environ Microbiol 80, 5884–5891 (2014). 10.1128/AEM.01763-14

46 Fisher, R. A., Gollan, B. & Helaine, S. Persistent bacterial infections and persister cells. Nat Rev Microbiol 15, 453–464 (2017). 10.1038/nrmicro.2017.42

47 Martin, M. Cutadapt removes adapter sequences from high-throughput sequencing reads. *EMBnet*. journal 17, 10–12 (2011).

48 Dobin, A. et al. STAR: ultrafast universal RNA-seq aligner. Bioinformatics 29, 15–21 (2013).

49 Howe, K. L. et al. Ensembl 2021. Nucleic acids research 49, D884–D891 (2021).

50 Langmead, B. & Salzberg, S. L. Fast gapped-read alignment with Bowtie 2. Nature methods 9, 357–359 (2012).

51 Li, H. et al. The sequence alignment/map format and SAMtools. bioinformatics 25, 2078–2079 (2009).

52 Liao, Y., Smyth, G. K. & Shi, W. featureCounts: an efficient general purpose program for assigning sequence reads to genomic features. Bioinformatics 30, 923–930 (2014).

53 Howe, K. L. et al. Ensembl Genomes 2020—enabling non-vertebrate genomic research. Nucleic acids research 48, D689–D695 (2020).

