## Supplementary Information for "Sample-derived cDNA guides broad host RNA depletion for *in vivo* pathogen transcriptomics"

#### Supplementary Note S1. Selection of SuperScript IV for cDNA synthesis in host RNA depletion experiments

To identify the most efficient reverse transcriptase for cDNA synthesis in the context of host RNA depletion, we compared five commercially available reverse transcriptases: RevertAid, RevertAid H(–), AffinityScript, SuperScript IV, and AMV. Equal amounts of total RNA were reverse transcribed under identical conditions using random hexamer priming, following each manufacturer's recommended protocol. Following cDNA synthesis, RNase H was added to digest the RNA template strand for enzymes lacking intrinsic RNase H activity, leaving only the newly synthesized single-stranded cDNA. The resulting cDNA products were resolved by gel electrophoresis and quantified by densitometry (**Supplementary Fig. 1a**). SuperScript IV yielded the highest cDNA signal (46.6 arbitrary units), substantially exceeding RevertAid H(–) (23.7), AffinityScript (23.4), RevertAid (19.7), and AMV (15.2) (**Supplementary Fig. 1b**). Based on these results, SuperScript IV was selected for all subsequent cDNA synthesis steps in the host RNA depletion protocol.

#### Supplementary Note S2. RT-qPCR validation of host-specific rRNA depletion

To further verify that our optimized cDNA-guided RNase H depletion does not deplete bacterial RNA, RT-qPCR was performed targeting host 18S rRNA and bacterial 16S rRNA before and after depletion across all experiments **(Supplementary Fig S5a)**. Depletion efficiency was calculated using the ΔCt method, comparing Ct values of depleted samples to non-depleted controls using the formula:

$$\text{Depletion (\%)} = (1 - 2^{-\Delta Ct}) \times 100$$

where $\Delta Ct = Ct_{(depleted)} - Ct_{(non\text{-}depleted)}$.

Across all experiments, cDNA-guided depletion consistently reduced host 18S rRNA levels by approximately 99%, while bacterial 16S rRNA levels remained unchanged, with no statistically significant reduction observed **(Supplementary Fig S5b)**. The preservation of bacterial RNA is mechanistically supported by the known dynamics of reverse transcription in mixed RNA templates. When random hexamers are used for cDNA synthesis in a host–bacterial RNA mixture, highly abundant host transcripts dominate and sequester the available hexamer primers, resulting in non-linear stoichiometry that strongly disfavors cDNA synthesis from rare bacterial transcripts[1-3]. Consequently, bacterial RNA:cDNA hybrids are rarely formed during RNase H digestion, leaving bacterial transcripts intact and available for downstream sequencing.

**Supplementary Note S3. Rationale for inclusion of bacterial-only RNA sequencing libraries**

To provide a reference baseline for evaluating the bacterial transcriptome composition in depleted and non-depleted host–bacterial RNA mixtures, *S. epidermidis* RNA isolated directly from bacterial culture was included as a bacterial-only control in RNA-seq experiments. These libraries were prepared from purified total bacterial RNA without any rRNA depletion step, preserving the native bacterial RNA composition including ribosomal RNA fractions. Since bacterial-only libraries contain no host RNA, sequencing reads map exclusively to the *S. epidermidis* genome, enabling full transcriptome coverage at substantially lower sequencing depths compared to host–bacterial mixture libraries. Bacterial-only libraries were therefore sequenced at lower depths than non-depleted and depleted mixture libraries, as rarefaction analysis confirmed that near-complete *S. epidermidis* gene detection was achieved at these depths **(Fig. 2e,f)**. These bacterial-only libraries served two purposes in the study. First, they were used as a transcriptional reference to assess whether cDNA-guided depletion distorts the global bacterial gene expression profile, by comparing bacterial transcript abundance and composition between depleted libraries and the bacterial-only baseline. Second, they were included in t-SNE embedding and differential expression analyses **(Fig. 5)** to establish whether depleted and non-depleted mixture libraries cluster with their expected transcriptional state rather than with stress-condition transcriptomes. Bacterial-only libraries were not included in the *in vivo* S. Typhimurium experiments, where the complexity and heterogeneity of infection samples precluded the use of a matched pure bacterial reference.

**Supplementary Note S4. Optimization and specificity of cDNA-guided host RNA depletion**

During optimization of the cDNA-guided depletion protocol, we compared two cDNA purification strategies: column-based purification and AMPure XP bead purification at 2× bead concentration. Bioanalyzer analysis of column-purified cDNA revealed a substantial population of small fragments (below ~100 nt) that were absent in bead-purified preparations **(Supplementary Fig. S7)**. To evaluate whether these small fragments affect depletion specificity, we performed RT-qPCR targeting host 18S rRNA and bacterial 16S rRNA following depletion with each cDNA preparation.

Depletion with bead-purified cDNA selectively reduced host 18S rRNA levels while preserving bacterial 16S rRNA abundance. In contrast, depletion with column-purified cDNA reduced both host 18S rRNA and bacterial 16S rRNA levels **(Supplementary Fig. S5)**, indicating non-specific hybridization of small cDNA fragments to bacterial RNA, which lack the sequence specificity required for selective host RNA targeting. Accordingly, AMPure XP bead purification at 2× bead concentration was adopted for all subsequent cDNA preparations.

**Supplementary Table S1. Oligonucleotide probe sequences for supplemental depletion of 7S RNA and 18S rRNA.** Sequences of oligonucleotide probes targeting regions of mouse and human 7S RNA and 18S rRNA. All sequences are listed in the 5' to 3' orientation.

| Probes | Sequence (5’ to 3’) |
|---|---|
| 7S-1 | CTCAGCCTCCCGAGTAGCTGGGACTACAGGCGTGCGCCACCGCGCCCGGC |
| 7S-2 | CCGAACTTAGTGCGGACACCCGATCGGCATAGCGCACTACAGCCCAGAAC |
| 7S-3 | TTCCGACCTGGGCCGGTTCACCCCTCCTTAGGCAACCTGGTGGTCCCCCG |
| 7S-4 | CCAGGCTGGAGTGCAGTGGCTATTCACAGGCGCGATCCCACTACTGATCA |
| 18S-1 | TGGTGCCCTTCCGTCAATTCCTTTAAGTTTCAGCTTTGCAACCATACTCC |
| 18S-2 | GTCTCGTTCGTTATCGGAATTAACCAGACAAATCGCTCCACCAACTAAGA |
| 18S-3 | GAACGCCACTTGTCCCTCTAAGAAGTTGGGGGACGCCGACCGCTCGGGGG |
| 18S-4 | AGTCAGTGTAGCGCGCGTGCAGCCCCGGACATCTAAGGGCATCACAGACC |
| 18S-5 | GGGCAGGGACTTAATCAACGCAAGCTTATGGCCCGCACTTACTGGGAATT |
| 18S-6 | CCGCAGGTTCACCTACGGAAACCTTGTTACGACTTTTACTTCCTCTAGAT |

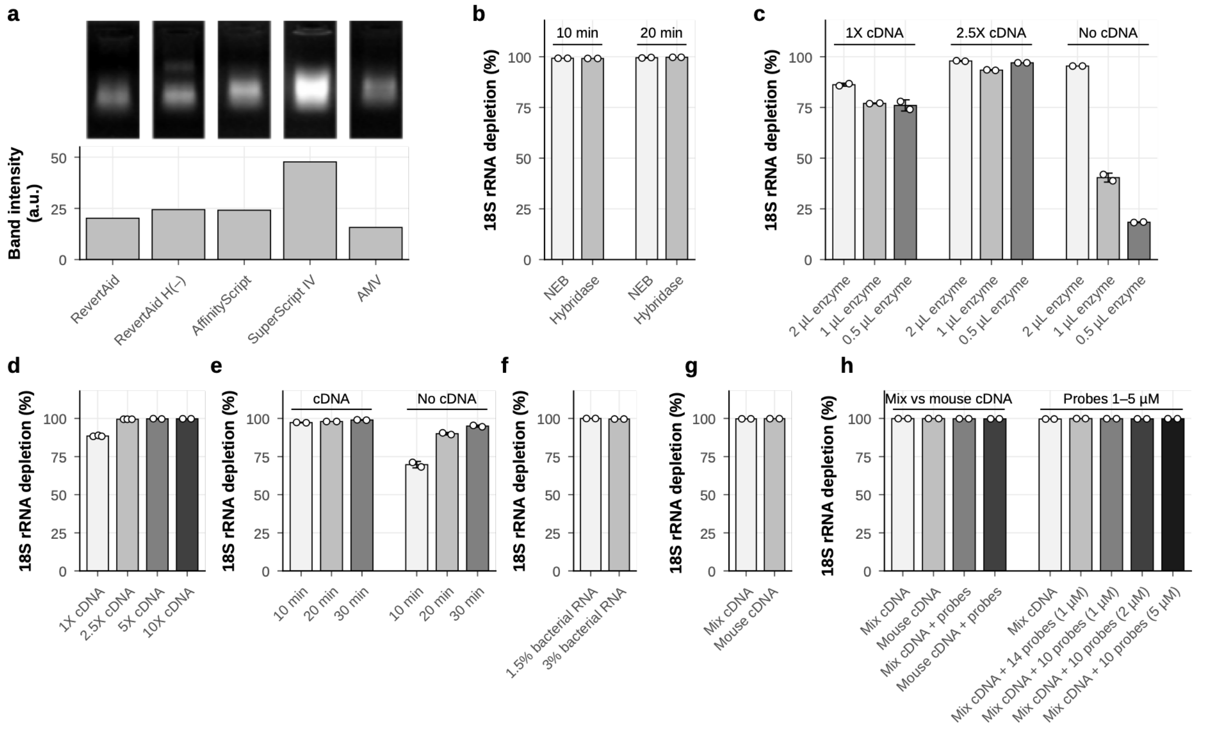


**Supplementary Fig. S1. Optimisation of cDNA-guided RNase H depletion. a** cDNA synthesis efficiency of five reverse transcriptases: RevertAid, RevertAid H(–), AffinityScript, SuperScript IV and AMV. Top, gel lanes showing cDNA remaining after RNase H digestion of the RNA template strand. Equal RNA input and equal loading volumes were used, and brightness is consistent across lanes. Bottom, mean background-subtracted pixel intensity of each lane (arbitrary units). **b–h** 18S rRNA depletion measured by RT-qPCR. **b** NEB or Hybridase RNAse H enzyme for 10 or 20 min. **c** 2, 1 or 0.5 µL of NEB RNAase H enzyme with 1X cDNA, 2.5X cDNA or no cDNA. **d** 1X, 2.5X, 5X or 10X cDNA used for depletion. **e** 10, 20 or 30 min of treatment with cDNA or with enzyme only (no cDNA). **f** Samples containing 1.5% or 3% bacterial RNA together with mouse cDNA. **g** Mix cDNA or mouse cDNA only. **h** Left, Mix cDNA or Mouse cDNA, with or without 18S rRNA and 7S RNA probes. Right, Mix cDNA alone or with 14 or 10 probes at 1–5 µM concentration. In **b**, **f** and **g**, each condition was compared with a matched no-cDNA control (same enzyme and incubation time in **b**; same bacterial RNA fraction in **f**). In **c** and **e**, all conditions were compared with a shared control containing no enzyme and no cDNA (n = 6). In d, the 1X/2.5X and 5X/10X conditions come from two separate experiments, each compared with its own no-cDNA control. In **h**, each group of bars is a separate experiment, and every condition is compared with the non-depleted sample from that experiment. Depletion was calculated as $(1 - 2^{-\Delta Cq}) \times 100$, where $\Delta Cq = Cq(condition) - Cq(control)$. Bars show the mean of technical replicates (n = 2; n = 3 for 1X and 2.5X in d). Dots show individual replicates, and error bars show s.d.

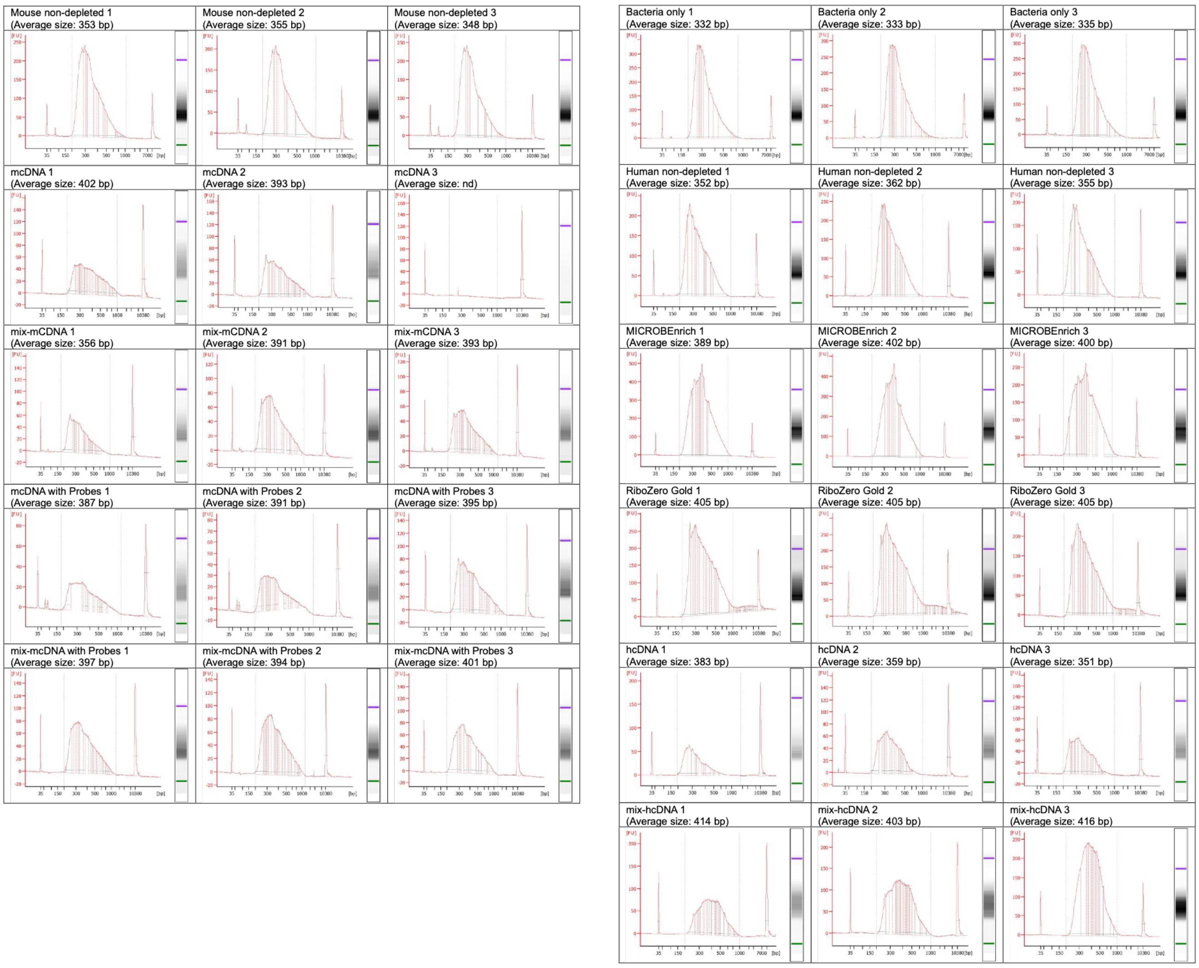


**Supplementary Fig. S2. Bioanalyzer size-distribution profiles of the sequencing libraries.** Electropherograms of the final libraries analysed on an Agilent 2100 Bioanalyzer with the High Sensitivity DNA assay. Each panel plots fluorescence intensity (FU) against fragment size (bp), with the corresponding virtual gel image shown on the right. The peaks at 35 bp and 10,380 bp are the lower (green) and upper (purple) size markers, respectively. Dashed vertical lines mark the region used to calculate the average library size, which is given above each panel. Libraries were prepared in triplicate. nd, not determined.

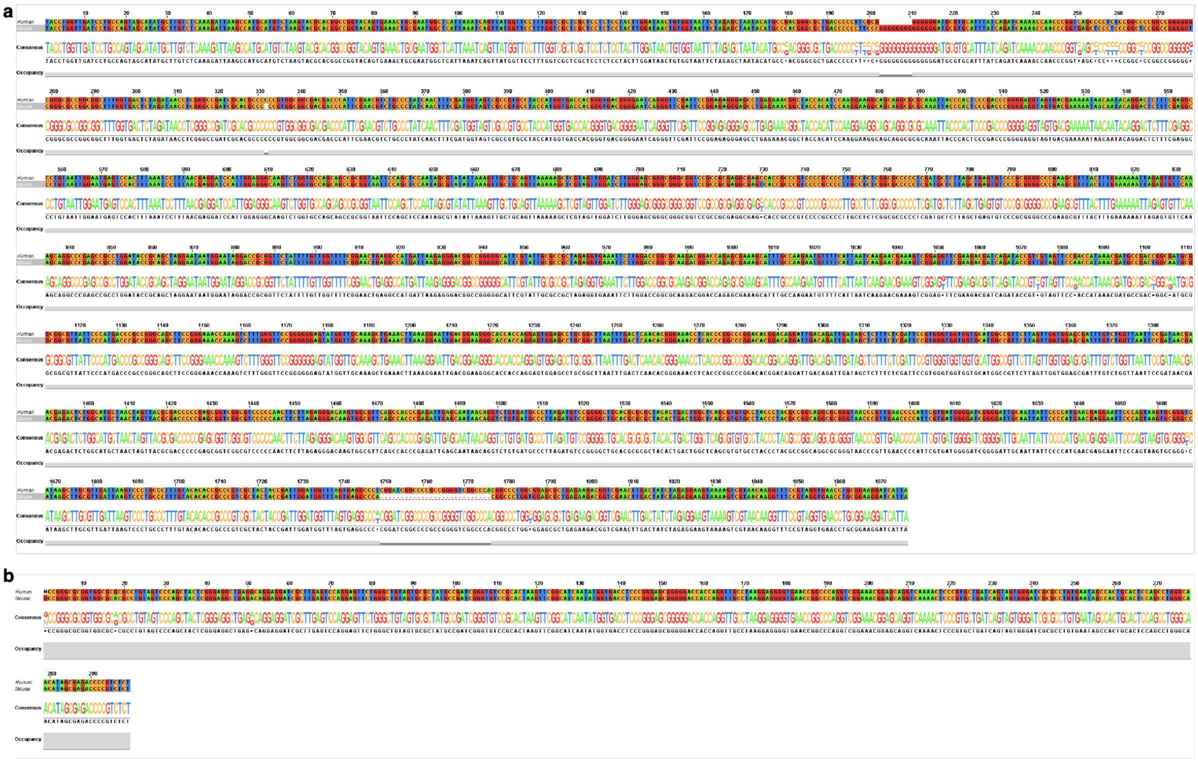


**Supplementary Fig. S3. Pairwise sequence alignment of human and mouse 18S and 7S RNA.** Pairwise alignment of human and mouse 18S rRNA (1,869 nt) **(a)** and 7S RNA (300 nt) **(b)** sequences. Nucleotides are color-coded by identity (A, red; T, blue; G, yellow; C, green), with conserved positions appearing as uniform columns across both sequences. Consensus and occupancy tracks are shown below each alignment block.

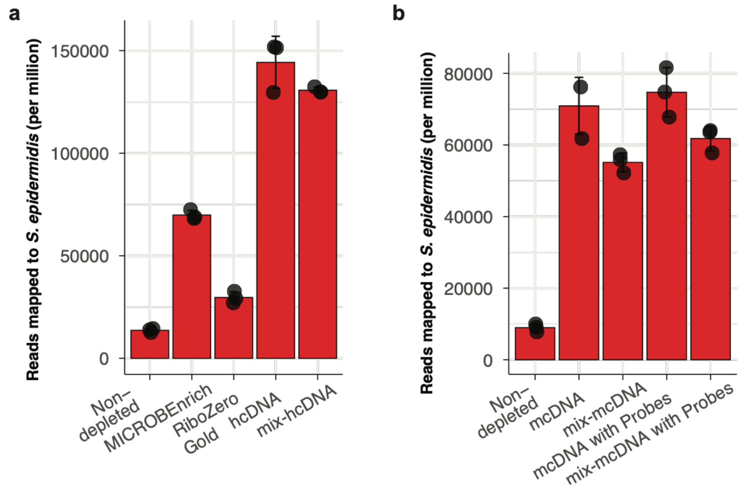


**Supplementary Fig. S4. Bacterial reads per million total reads across depletion conditions in human and mouse RNA mixtures.** Bar plots showing the number of reads mapping to the *S. epidermidis* genome per million total reads for human **(a)** and mouse **(b)** RNA mixtures across all depletion conditions. In human mixtures **(a)**, cDNA-guided depletion with hcDNA and mix-hcDNA yielded ~145,000 and ~132,000 reads per million, respectively, compared to MICROBEnrich (~70,000), RiboZero Gold (~28,000), and non-depleted samples (~15,000). In mouse mixtures **(b)**, mcDNA with probes and mix-mcDNA with probes achieved ~75,000 and ~65,000 reads per million, respectively, compared to non-depleted samples (~10,000). Data represent mean ± SEM; individual replicates shown as black dots.

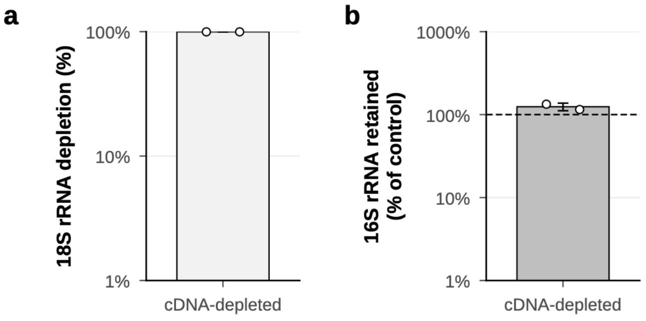


**Supplementary Fig. S5. cDNA-guided depletion removes host 18S rRNA while preserving bacterial 16S rRNA.** Host 18S rRNA and bacterial 16S rRNA were quantified by RT-qPCR after depletion mouse cDNA, relative to a non-depleted sample processed in parallel. a 18S rRNA depletion (log scale). b 16S rRNA retained (log scale); the dashed line marks the non-depleted level. Bars show the mean of technical replicates, dots show individual replicates, and error bars show s.d.

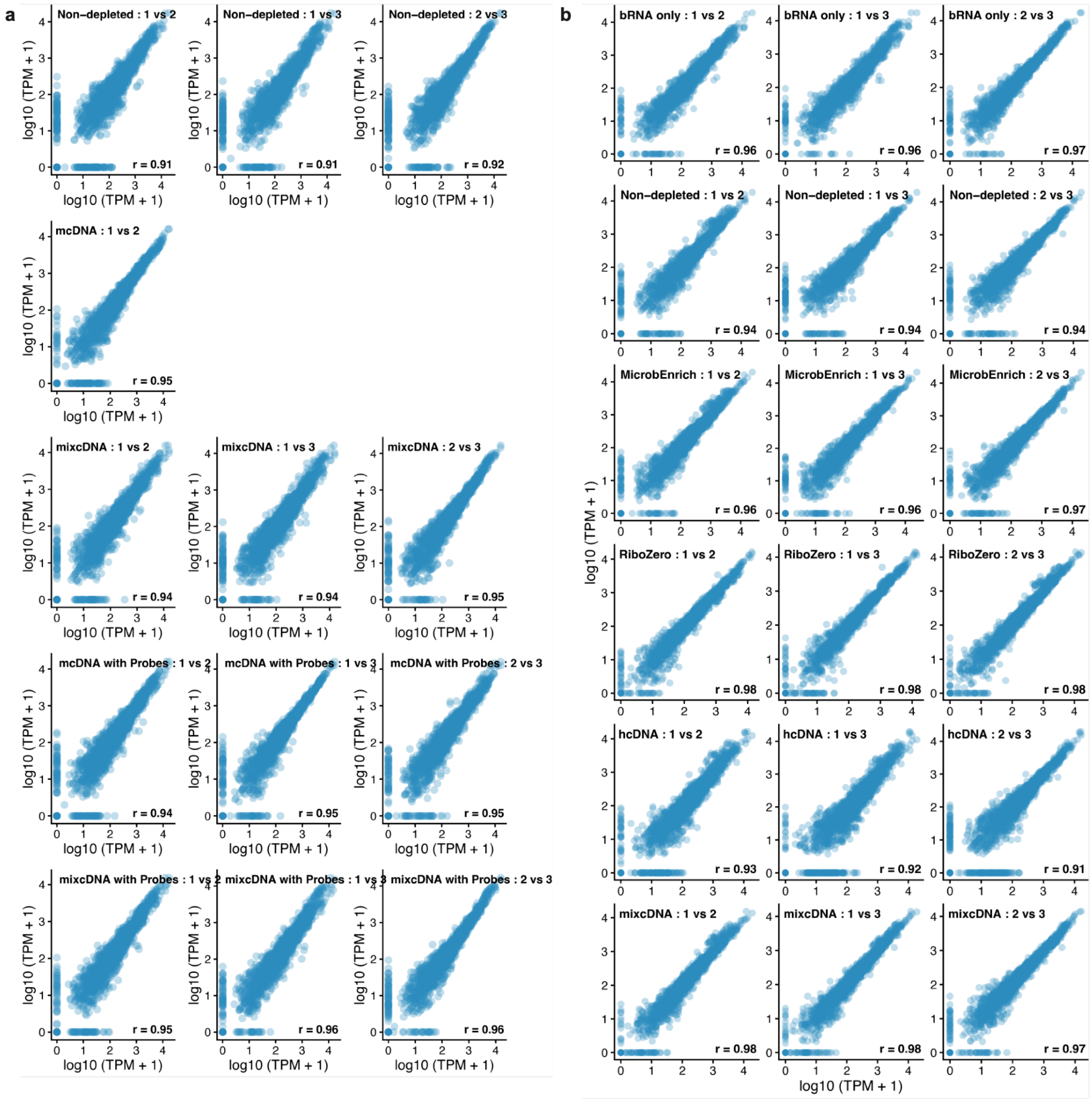


**Supplementary Fig. S6. Pairwise scatterplot comparison of bacterial transcriptome profiles.** Gene-level comparisons of *S. epidermidis* transcript abundance between biological replicates (1 vs 2, 1 vs 3, and 2 vs 3) across RNA preparation, depletion, and enrichment conditions for mouse **(a)** and human **(b)** RNA mixtures. Axes represent transcript abundances as $\log_{10}$(TPM+1). Pearson correlation coefficient (r) is displayed for each subplot.

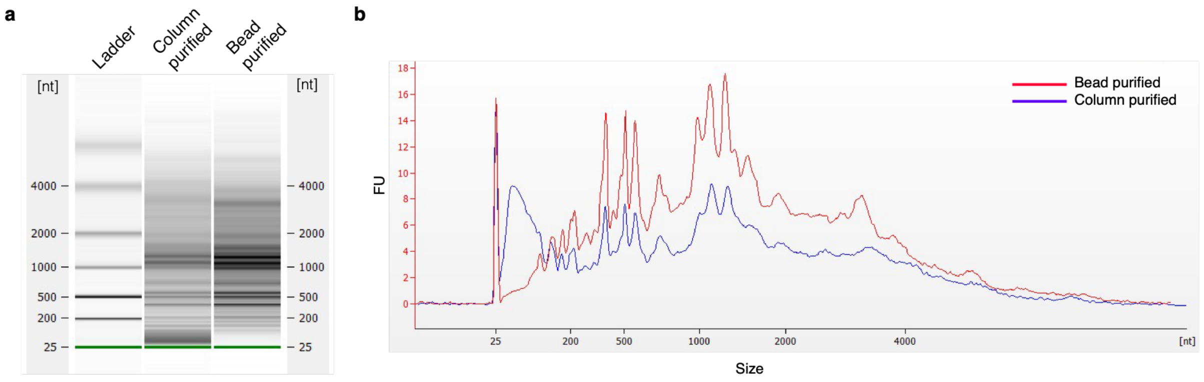


**Supplementary Fig. S7. Bioanalyzer electropherogram profiles of purified cDNA. (a)** Gel image and **(b)** electropherogram traces showing cDNA fragment size distribution following column (Zymo) purification (column purified, blue) and 2× AMPure bead (Beckman Coulter) purification (bead purified, red). The x-account represents fragment size (nucleotides) and the y-axis represents fluorescence units (FU).

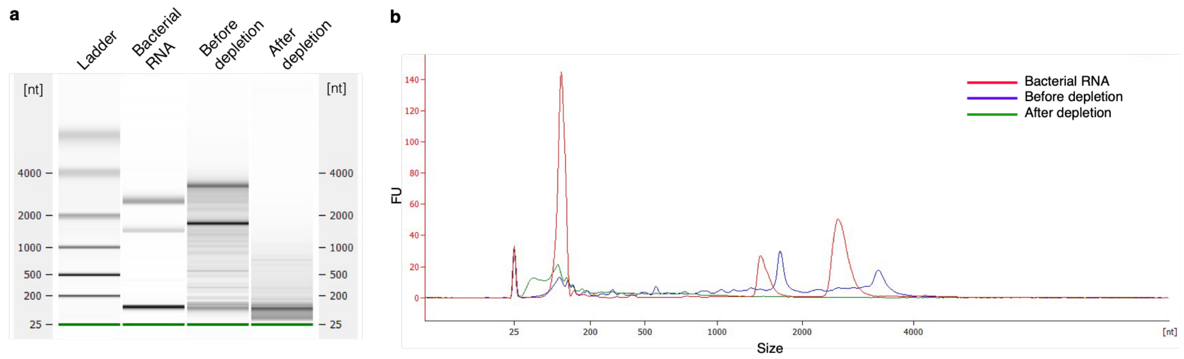


**Supplementary Fig. S8. Bioanalyzer electropherogram profiles of RNA samples before and after cDNA-guided depletion. (a)** Gel image and **(b)** electropherogram traces showing RNA fragment size distribution for purified *S. epidermidis* total RNA (Bacterial RNA, red), mixed RNA containing 10% bacterial and 90% mouse total RNA prior to depletion (Before depletion, blue), and mixed RNA following cDNA-guided host RNA depletion (After depletion, green). The x-axis represents fragment size (nucleotides) and the y-axis represents fluorescence units (FU).

### References

1 Moro, G., Brunner, E. & Basler, K. A practical guide to targeted single-cell RNA sequencing technologies. *Communications Biology* **9**, 250 (2026). https://doi.org/10.1038/s42003-026-09675-y

2 Minshall, N. & Git, A. Enzyme- and gene-specific biases in reverse transcription of RNA raise concerns for evaluating gene expression. *Sci Rep* **10**, 8151 (2020). https://doi.org/10.1038/s41598-020-65005-0

3 Verwilt, J., Mestdagh, P. & Vandesompele, J. Artifacts and biases of the reverse transcription reaction in RNA sequencing. *Rna* **29**, 889-897 (2023).
